# Notch1 cortical signaling regulates epithelial architecture and cell-cell adhesion

**DOI:** 10.1101/2023.01.23.524428

**Authors:** Matthew J. White, Kyle A. Jacobs, Tania Singh, Matthew L. Kutys

## Abstract

Notch receptors control tissue morphogenic processes that involve coordinated changes in cell architecture and gene expression, but how a single receptor can produce these diverse biological outputs is unclear. Here we employ a 3D organotypic model of a ductal epithelium to reveal tissue morphogenic defects result from loss of Notch1, but not Notch1 transcriptional signaling. Instead, defects in duct morphogenesis are driven by dysregulated epithelial cell architecture and mitogenic signaling which result from loss of a transcription-independent Notch1 cortical signaling mechanism that ultimately functions to stabilize adherens junctions and cortical actin. We identify that Notch1 localization and cortical signaling are tied to apical-basal cell restructuring and discover a Notch1-FAM83H interaction underlies stabilization of adherens junctions and cortical actin. Together, these results offer new insights into Notch1 signaling and regulation, and advance a paradigm in which transcriptional and cell adhesive programs might be coordinated by a single receptor.

## INTRODUCTION

Coordinated changes in cell-cell adhesions are central to tissue morphogenesis and integrity, and thus are essential mediators in development, tissue physiology, and disease pathogenesis (Belardi et al., 2020; Borghi and Nelson, 2009). Cell-cell adhesive junctions coordinate multicellular behavior by transmitting mechanical forces between cells and by influencing the localization, duration, and cytoskeletal coupling of receptor interactions to orchestrate juxtracrine signaling (Collinet and Lecuit, 2021; Gumbiner, 1996). Yet, at a fundamental level, molecular mechanisms that orchestrate cell-cell junction assembly and stability are not well understood. Similarly, it remains unclear how junctional changes are synchronized with transcriptional programs, for example in the context of the coupled patterning of cell lineages and movement during developmental morphogenesis (de Celis et al., 1996; Falo-Sanjuan and Bray, 2022).

The Notch family of receptors is a fundamental, conserved regulator of developmental patterning, where receptor signaling determines cell fates and patterns of gene expression amongst neighboring cells (Bray, 2016; Siebel and Lendahl, 2017). Notch receptors are integrated into the plasma membrane as noncovalent heterodimers composed of a large extracellular domain (ECD) polypeptide bound to a transmembrane fragment that consists of an extracellular sequence and a transmembrane domain (hereafter collectively referred to as the transmembrane domain (TMD)), and an intracellular domain (ICD). Notch receptors are activated via interaction with ligands presented on adjacent cells, which occurs through multiple steps that are independently gated by sequential events. Mechanical force applied to the Notch ECD causes conformational unfolding of an extracellular negative regulatory region that renders the receptor sensitive to sequential proteolytic cleavages, first at an extracellular S2 site removing the ECD and subsequently at an intramembrane S3 site by the γ-secretase complex cleaving the ICD from the TMD (Gordon et al., 2008; Gordon et al., 2015; Kopan and Ilagan, 2009; Kovall et al., 2017). Notch exerts transcriptional effects through cleaved ICD, which can translocate to the nucleus and form a Notch transcription activation complex with cofactors RBPJ and MAML1/2 (Borggrefe and Oswald, 2009; Wang et al., 2014). This Notch signaling mechanism is conserved across the entire animal kingdom, and Notch activation is implicated during tissue morphogenesis in many developmental programs (Hellstrom et al., 2007; Lloyd-Lewis et al., 2019; Priya et al., 2020). However, how morphogenetic changes, which require dynamic cell architectures and cell-cell adhesions, may be regulated in the context of key developmental pathways like Notch remains largely unaddressed.

The Notch gene was discovered from a mutant allele in *Drosophila* causing the formation of a wing “notch” due to defects in the dorsoventral compartmentalization of the developing wing disc epithelium. Cells along the dorsoventral boundary have distinctive properties and their specification requires Notch activity (de Celis et al., 1996). Notch-dependent, distinctive filamentous actin, myosin II, and adherens junction phenotypes also form at the dorsoventral boundary, yet these changes are not accounted for purely by the transcriptional regulation of target genes associated with Notch activation (Major and Irvine, 2005, 2006). In mammals, Notch1 is expressed broadly in adult epithelium as well as endothelium, and evidence suggest interactions with adherens junctions and actomyosin are linked to both Notch receptor activation and downstream function in distinct contexts (Crowner et al., 2003; Falo-Sanjuan and Bray, 2021; Hunter et al., 2019; Khait et al., 2016; Kwak et al., 2022; Lowell and Watt, 2001; Priya et al., 2020). Further, loss of Notch1 impairs epithelial and endothelial monolayer barrier function, a tissue property that is dependent upon stable cell–cell adhesion (Blanpain et al., 2006; Dahan et al., 2011; Demehri et al., 2008; Movahedan et al., 2013; Polacheck et al., 2017). We recently described a mechanism by which hemodynamic forces of blood flow activate the Notch1 receptor in the vascular endothelium to enhance vessel barrier function. This process does not involve ICD-mediated transcription, but instead operates through a mechanism we refer to as Notch1 cortical signaling, where the TMD acts as a focal point of protein-protein interactions in a pathway that enhances endothelial adherens junctions (Polacheck et al., 2017). If, and how, transcription-independent Notch1 cortical signaling influences epithelial cell architecture and cell-cell adhesion is unexplored.

In the present study, we employ an organotypic microfluidic platform capable of recapitulating and dissecting three-dimensional (3D) morphogenic features of a ductal epithelium. Using CRISPR– Cas9 gene editing to specifically isolate Notch1 cortical or transcriptional signaling in human epithelia, we observe distinct 3D morphogenic consequences in a ductal epithelium result upon loss of Notch1 cortical signaling, but not transcriptional signaling. Tissue morphogenic defects are driven by dysregulated epithelial cell architecture and mitogenic signaling, which Notch1 cortical signaling controls through the stabilization of adherens junctions and cortical actin organization. Mechanistically, we find that Notch1 receptor localization and cortical signaling function are tied to epithelial apical-basal cell columnar restructuring, and we unbiasedly identify molecular interactions that underlie adherens junction regulation by Notch1 cortical signaling.

## RESULTS

### Notch1 influences ductal epithelium morphogenesis through a transcription-independent mechanism

To specifically isolate Notch1 signaling functions independent of the ICD and transcription, we applied methods that we previously established to generate endogenous Notch1 truncation mutants in primary human endothelia (Polacheck et al., 2017). We engineered human mammary epithelial cells (MCF10A) harboring either CRISPR/Cas9-mediated ablation of Notch1 (*NOTCH1^KO^*) or truncation of the Notch1 intracellular domain (*ICD^KO^*) which preserves the ECD and TMD (Figure 1A). Cell lysis, heterodimer dissociation via SDS-PAGE, and western blotting permits visualization of Notch1 ECD or transmembrane fragment polypeptides at distinct molecular weights using respective antibodies. Western blot confirmed deletion of Notch1 in *NOTCH1^KO^* epithelia, as well as ICD truncation of the transmembrane fragment and preservation of the ECD in *ICD^KO^* epithelia. *NOTCH1^KO^* or *ICD^KO^* epithelia do not have altered E-cadherin protein levels or phosphorylation of β-catenin compared to a non-targeting scramble guide RNA (SCR) control (Figure 1A).

**Figure 1.**
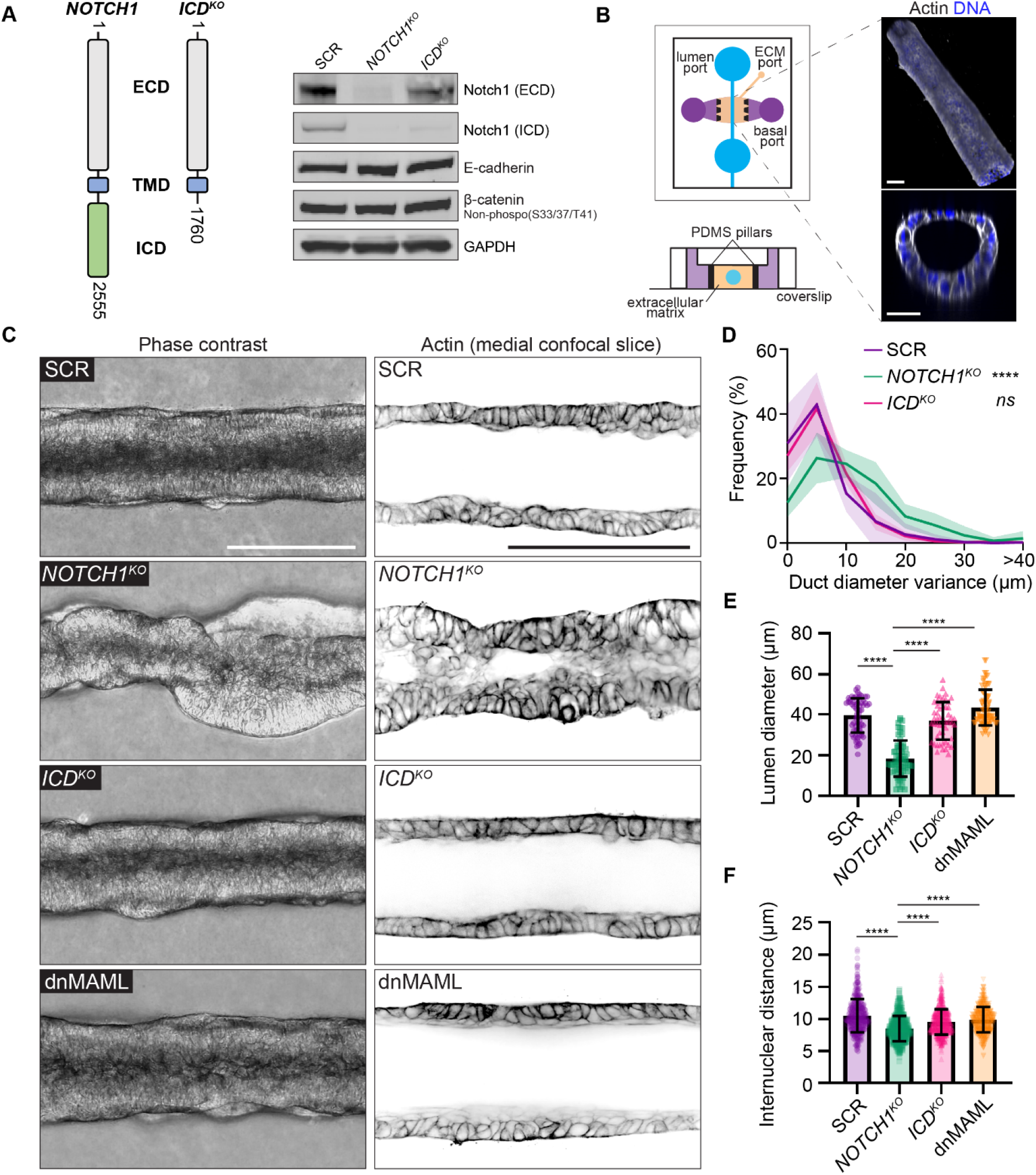
Notch1 influences ductal epithelium architecture through a transcription-independent mechanism. (A) Left: Schematic of endogenous Notch1 CRISPR-Cas9 mutant used to truncate Notch1 ICD. Right: Western blot of lysates from scramble control (SCR), *NOTCH1^KO^*, and *ICD^KO^* monolayers immunoblotted for Notch1 ECD, Notch1 ICD, E-Cadherin, β-Catenin (nonphosphorylated (S33/37/T41)), and GAPDH. (B) Left: Organotypic microfluidic platform consisting of an engineered 3D mammary ductal epithelium embedded in physiologic ECM. Luminal ports (blue) used for cell seeding and perfusion of medium through the lumen, basal ports (purple) used for delivery of medium containing growth factors, ECM injection port and ECM compartment (beige), PDMS pillars (black) used to contain hydrogel ECM (Top schematic: top-down view; bottom schematic: cross-section). Right: Representative 3D oblique projection (top) and cross-section (bottom) of a 3D mammary duct labeled with phalloidin (grey) and Hoechst (blue). Scale bars, 50 μm. (C) Left: Representative phase contrast micrographs of SCR, *NOTCH1^KO^, ICD^KO^*, and dnMAML ducts. Scale bar, 150 μm. Right: Representative medial confocal slice fluorescence micrographs of SCR, *NOTCH1^KO^, ICD^KO^*, and dnMAML epithelial ducts labeled with phalloidin (black). Scale bar, 100 μm. (D) Quantification of duct diameter variance measured from phase contrast micrographs as shown in C. n = 8, 7, 7 ducts from three independent experiments. (E) Quantification of lumen diameter measured from confocal micrographs of phalloidin as shown in C. n ≥ 10 average lumen diameters, from three independent experiments. (F) Quantification of internuclear distances measured from confocal micrographs of Hoechst labeled ducts. n ≥ 40 internuclear distances per duct from three independent experiments. For all plots, mean ± SEM; one-way ANOVA with Tukey’s post-hoc test, ****p<0.0001.

We first investigated whether Notch1 influences the assembly or maintenance of a 3D organotypic human mammary ductal epithelium. To maintain control over cells, extracellular matrix (ECM), and tissue architecture, we developed a microfluidic model of a 3D mammary ductal epithelium which consists of a channel surrounded by ECM that is lined with human mammary epithelial cells and supported by basal delivery of growth factors (Figure 1B). Seeded SCR cells quickly populate the surface of the duct channel and form stable, non-invasive monolayers over the course of seven days. The resulting linear ductal tissues have a hollow central lumen that is lined by columnar epithelial cells that are growth arrested. Seeded *NOTCH1^KO^* cells similarly populate the channel surface, but in contrast develop tortuous duct architectures with prominent tissue outgrowths which cause large variance in duct diameter (Figure 1C,D; Supplemental Figure 1A). Perfusion of duct lumens with fluorescent microbeads (4 μm) indicated that *NOTCH1^KO^* lumens are occluded compared to SCR ducts (Supplementary Figure 1B). Further, medial confocal sections of *NOTCH1^KO^* ducts revealed a failure to form an ordered monolayer and extensive lumen cell in-filling, a two-fold decrease in average lumen diameter (Figure 1C,E), and increased cell packing as quantified by reduced internuclear distances (Figure 1C,F; Supplemental Figure 1C).

To investigate whether transcription-independent functions of Notch1 contribute to the *NOTCH1^KO^* duct phenotype, we generated ducts from *ICD^KO^* cells or cells engineered to express a dominant-negative form of the Notch transcriptional cofactor mastermind-like protein 1 (dnMAML) (Polacheck et al., 2017; Weng et al., 2003). dnMAML expression decreases transcript levels of Notch transcriptional target genes HES1 and HEY1 (Supplemental Figure 1D). *ICD^KO^* or dnMAML ducts closely resemble the overall tissue architecture of SCR ducts, with slightly larger lumen diameters forming in dnMAML ducts (Figure 1C-E). Importantly, no evidence of lumen cell infilling was present in either *ICD^KO^* or dnMAML ducts (Figure 1C-F). Taken together, *NOTCH1^KO^* cells form 3D ducts with tortuous architectures, tissue outgrowths, and occluded lumens, and these phenotypes are not present in ducts constructed from cells harboring two distinct perturbations that suppress Notch1 transcription. This identifies a specific function for transcription-independent Notch1 cortical signaling in regulating the assembly or maintenance of a 3D mammary ductal epithelium.

### Aberrant cell architecture and proliferation underlie *NOTCH1^KO^* duct defects

We next investigated which cell behaviors contribute to the *NOTCH1^KO^* duct phenotype through a temporal analysis of duct assembly. SCR and *NOTCH1^KO^* cells similarly adhere to the channel architecture and progressively form a monolayer. Approximately three days after initial seeding, SCR ducts initiate expansion of duct lumens; however, lumen expansion is significantly diminished in *NOTCH1^KO^* ducts and small tissue outgrowths are detectable by phase contrast microscopy (Supplementary Figure 1A). Using a cell-permeable fluorescent probe for filamentous actin and live confocal microscopy, we visualized cell dynamics within *NOTCH1^KO^* ducts at the onset of these morphogenic differences. Timelapse imaging revealed several areas of cell multilayering that are formed from frequent cell divisions oriented orthogonal to the basal ECM interface. Daughter cells positioned inward amassed in the lumen and typically did not reintegrate into the duct monolayer lining (Figure 2A, Supplementary Video 1). Consistent with observations from 3D mammary epithelial acinar models (Jaffe et al., 2008), SCR and *ICD^KO^* cells instead orient spindle axes along the basal ECM interface during division (Supplementary Figure 2A). Pulse labeling ducts with 5-ethynyl-2’-deoxyuridine (EdU) to assess cell proliferation revealed increased EdU incorporation in *NOTCH1^KO^* ducts relative to SCR or *ICD^KO^* that is primarily localized within cell masses in the lumen (Figure 2B, D). These observations indicated dysregulated epithelial architecture and proliferation may be associated with loss of Notch1 cortical signaling in *NOTCH1^KO^* ducts.

**Figure 2.**
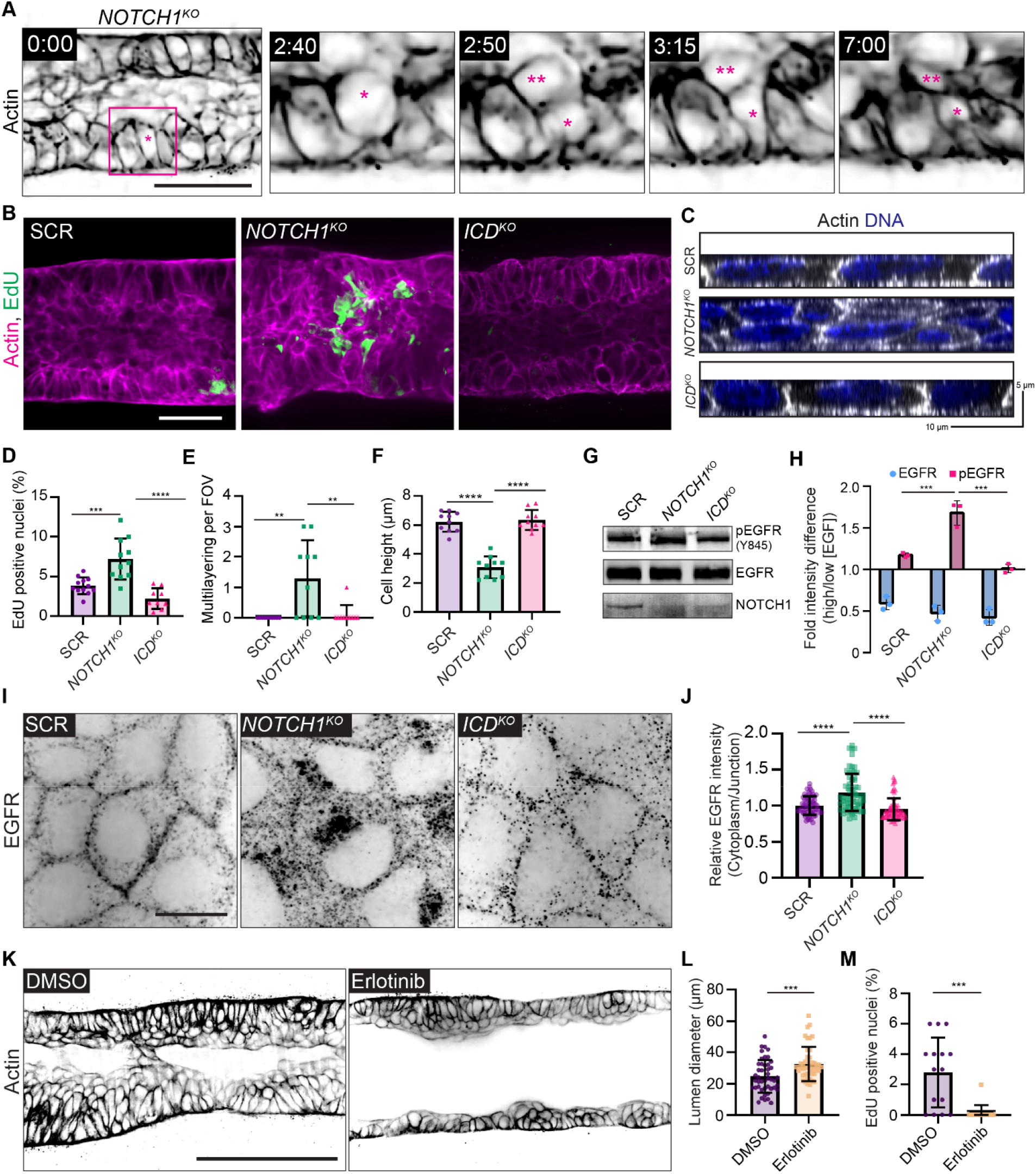
Notch1 cortical signaling regulates epithelial cell architecture and suppresses EGFR mitogenic signaling. (A) Individual time frames from a live cell movie of actin within a *NOTCH1^KO^* duct labeled with SPY650-FastAct (black). Inset for individual time frames outlined in magenta in the 0:00-hour frame. Parent cell is labeled with (*) and daughter cell is labeled with (**). Scale bar, 50 μm. (B) Maximum projection micrographs of 30 μm medial stacks of SCR, *NOTCH1^KO^*, and *ICD^KO^* epithelial ducts labeled with phalloidin (magenta) and EdU (green). Scale bar, 30 μm. (C) YZ orthogonal projections from fluorescence micrographs of SCR, *NOTCH1^KO^*, and *ICD^KO^* monolayers labeled with phalloidin (grey) and Hoechst (blue). (D) Quantification of the percentage of EdU positive nuclei in ducts. n ≥ 9 ducts from three independent experiments. (E) Quantification of regions of cell multilayering per field of view in fluorescence micrographs of SCR, *NOTCH1^KO^*, and *ICD^KO^* monolayers. n ≥ 10 fields of view from three independent experiments. (F) Quantification of cell height from SCR, *NOTCH1^KO^*, and *ICD^KO^* monolayers plated on hydrogels. n ≥ 10 fields of view from three independent experiments. (G) Western blot of lysates from confluent SCR, *NOTCH1^KO^*, and *ICD^KO^* monolayers immunoblotted for pEGFR (Y845), EGFR, and Notch1. (H) Quantification of western blot intensity difference of pEGFR and total EGFR levels in monolayers stimulated with EGF. n = 3 independent experiments. (I) Fluorescence micrographs of SCR, *NOTCH1^KO^*, and *ICD^KO^* monolayers immunostained for EGFR (black). Scale bar, 10 μm. (J) Quantification of relative cytoplasmic to junctional EGFR intensity. *n* ≥ 30 cells from three independent experiments. (K) Representative medial confocal slice micrographs of *NOTCH1^KO^* ducts treated with DMSO or 1 μM Erlotinib labeled with phalloidin (black). Scale bar, 100 μm. (L) Quantification of duct lumen diameter. n ≥ 15 average diameters from three independent experiments. (M) Quantification of the percentage of EdU positive nuclei in *NOTCH1^KO^* monolayers treated with DMSO or 1 μM Erlotinib. n ≥ 15 fields of view from three independent experiments. For plots D-F, H, and J, mean ± SEM; one-way ANOVA with Tukey’s post-hoc test, **p<0.01, ***p<0.001, ****p<0.0001. For plots L and M, mean ± SEM; twotailed unpaired t test, ***p<0.001.

Following these observations, we next modeled the physiological stiffness of basement membrane and underlying ECM by plating cells on compliant composite hydrogels in an effort to recapitulate cell morphodynamics observed in our 3D organotypic model (Nyga et al., 2021). Consistent with increased proliferation observed in ducts, *NOTCH1^KO^* cells cultured on compliant hydrogels display elevated EdU labeling compared to SCR, *ICD^KO^*, and dnMAML cells (Supplemental Figure 2B, C). Analysis of monolayer organization in the z-plane orthogonal to the substrate revealed that only *NOTCH1^KO^* monolayers contain regions of cell multilayering that are reminiscent of 3D duct cell lumen in-growth. In regions lacking multilayering, *NOTCH1^KO^* epithelia are two-fold shorter with diminished columnar cell morphology compared to SCR and *ICD^KO^* monolayers (Figure 2C,E-F; Supplemental Figure 2D). Decreased cell height and impaired columnar morphology, and elevated proliferation are similarly observed in human intestinal and bronchial *NOTCH1^KO^* monolayers relative to control (Supplemental Figure 3A-D). Thus, loss of Notch1 cortical signaling results in elevated proliferation, cell multilayering, and impaired apical-basal cell architecture on 2D hydrogels and within 3D organotypic ducts.

### Notch1 cortical signaling suppresses EGFR phosphorylation, internalization, and mitogenic signaling

To begin to characterize this Notch1 cortical signaling mechanism, we first focused on identifying molecular pathways leading to elevated epithelial proliferation. Notch signaling interacts with the Hippo/YAP growth control pathway (Totaro et al., 2017), but there are no differences in nuclear YAP localization between SCR and *NOTCH1^KO^* monolayers cultured on compliant hydrogels (Supplementary Figure 4A). The receptor tyrosine kinase epidermal growth factor receptor (EGFR) is a critical regulator of mammary tissue expansion during development and adult life, so we hypothesized that EGFR activity may be negatively regulated by Notch1 cortical signaling. *NOTCH1^KO^* monolayers cultured in high EGF containing medium (20 ng/ml) have elevated levels of active, tyrosine phosphorylated EGFR (Y845) compared to SCR and *ICD^KO^* (Figure 2G). Given this difference in EGFR phosphorylation, we next examined EGFR phosphorylation dynamics in response to EGF ligand. Comparing EGFR activity levels in monolayers cultured in low EGF containing medium (2 ng/ml) to those stimulated with medium containing high EGF (20 ng/ml) revealed a 1.7-fold increase in pEGFR in *NOTCH1^KO^* monolayers (Figure 2H). This suggests that Notch1 cortical signaling negatively regulates EGF sensitivity and EGFR phosphorylation in mammary epithelial cells.

Localization to adherens junctions negatively regulates EGFR activity, while internalization of active EGFR can enable mitogenic signaling (Lemmon and Schlessinger, 2010; Sullivan et al., 2022). Immunoprecipitation of E-cadherin from SCR and *NOTCH1^KO^* monolayer lysates shows a substantial decrease in E-cadherin association with EGFR upon deletion of Notch1 (Supplementary Figure 4C). Indeed, EGFR localizes to cell-cell interfaces in SCR and *ICD^KO^* monolayers; however, *NOTCH1^KO^* monolayers have diminished EGFR cell-cell contact localization and increased internalized cytoplasmic localization (Figure 2I,J; Supplementary Figure 4B). To causally relate EGFR kinase activity to the *NOTCH1^KO^* duct morphogenic defect, ducts were treated with the EGFR kinase inhibitor Erlotinib at three days post seeding. Erlotinib treatment significantly reduces *NOTCH1^KO^* lumen cell in-filling and proliferation, increases lumen sizes, but did not fully ameliorate disordered cell multilayering compared to vehicle control (Figure 2K-M). Altogether, these data support a model in which elevated EGFR kinase activity contributes to the aberrant proliferation observed upon loss of Notch1 cortical signaling, but EGFR activity is not responsible for defects in epithelial architecture and monolayer organization.

### Notch1 cortical signaling stabilizes adherens junctions and cortical actin

Adherens junctions and associated actomyosin networks are critical regulators of cell-cell contact dependent growth regulation. One mechanism by which this occurs is through suppression of EGFR mobility, internalization, and mitogenic signaling by stable adherens junctions (Chiasson-MacKenzie et al., 2015; Qian et al., 2004). Further, tension at adherens junctions is necessary for accurate orientation of epithelial cell division (Lisica et al., 2022). Our previous work identified that Notch1 cortical signaling strengthens vascular endothelial adherens junctions in response to hemodynamic shear stress (Polacheck et al., 2017). We therefore hypothesized loss of Notch1 cortical signaling causes defects in EGFR-driven proliferation and cell architecture through the alteration of adherens junctions.

Examination of SCR, *NOTCH1^KO^*, and *ICD^KO^* monolayers cultured on compliant ECM hydrogels revealed differences in the organization of E-cadherin-based adherens junctions specifically within *NOTCH1^KO^* monolayers. SCR and *ICD^KO^* adherens junctions are overall linear and continuous, however *NOTCH1^KO^* adherens junctions are discontinuous and oriented orthogonal to the cell-cell interface (Figure 3A, Supplementary Figure 5A). This distinct junction morphology is reminiscent of focal adherens junctions, which are immature adherens junctions that are typically associated with radially oriented actin fibers and posited to bear increased tension (Figure 3A,B) (Oldenburg et al., 2015). Indeed, while cortical actin is tightly enriched at SCR and *ICD^KO^* cell-cell interfaces, *NOTCH1^KO^* actin fibers are less cortically compact and fail to align parallel to the cell-cell interface (Figure 3C,D; Supplemental Figure 5B). E-cadherin-based adherens junctions and cortical actin organization are similarly disordered in *NOTCH1^KO^* human intestinal and bronchial epithelia (Supplemental Figure 3A,B). Increased cell-ECM traction forces are associated with destabilized adherens junctions in epithelial monolayers (Mertz et al., 2013; Scarpa et al., 2015). Traction force microscopy identified a 1.5-fold increase in relative cell-substrate tractions in *NOTCH1^KO^* monolayers relative to SCR or *ICD^KO^* (Figure 3E, F), further indicating destabilization of adherens junctions specifically upon loss of Notch1 cortical signaling.

**Figure 3.**
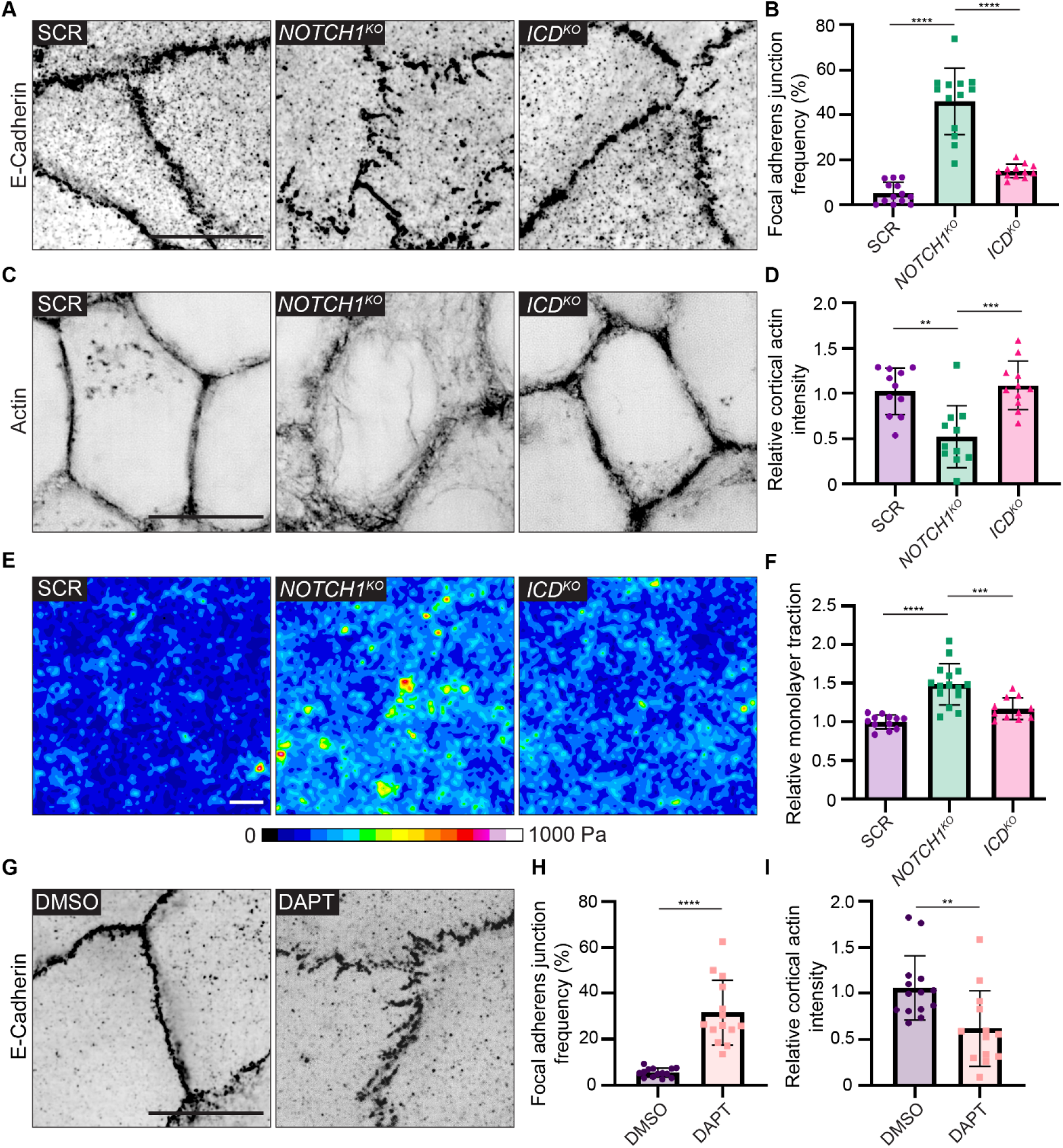
Notch1 cortical signaling stabilizes adherens junctions and cortical actin organization. (A) Super resolution by optical pixel reassignment (SoRa) fluorescence micrographs SCR, *NOTCH1^KO^*, and *ICD^KO^* monolayers immunostained for E-Cadherin (black). Scale bar, 10 μm. (B) Quantification of the frequency of focal adherens junctions. n ≥ 12 fields of view from three independent experiments. (C) SoRa fluorescence micrographs of SCR, *NOTCH1^KO^*, and *ICD^KO^* monolayers labeled with phalloidin (black). Scale bar, 10 μm. (D) Intensity of cortical actin at cell–cell junctions, quantified from phalloidin-stained micrographs. n ≥ 12 fields of view from three independent experiments. (E) Traction force microscopy traction maps averaged from ten fields of view from SCR, *NOTCH1^KO^*, and *ICD^KO^* monolayers. Scale bar, 20 μm. (F) Quantification of relative integrated monolayer tractions. n ≥ 11 traction force measurements from three independent experiments. (G) Fluorescence micrographs of wild-type cells treated with DMSO or 10 μM DAPT for two hours and immunostained with E-Cadherin (black). Scale bar, 10 μm. (H) Quantification of the frequency of focal adherens junctions in DMSO and DAPT treated cells. n ≥ 13 fields of view from three independent experiments. (I) Intensity of cortical actin at cell-cell junctions, quantified from phalloidin-stained micrographs of DMSO and DAPT treated cells. n ≥ 13 fields of view from three independent experiments. For plots B, D, and F, mean ± SEM; one-way ANOVA with Tukey’s post-hoc test, **p<0.01, ***p<0.001, ****p<0.0001. For plots H and I, mean ± SEM; two-tailed unpaired t test, **p<0.01, ****p<0.0001.

Our previous work identified ICD cleavage from the transmembrane fragment is necessary for the stabilization of endothelial adherens junctions by Notch1 cortical signaling (Polacheck et al., 2017). We therefore acutely treated wild type epithelial monolayers with DAPT (*N*-[*N*-(3,5-difluorophenacetyl)-L-alanyl]-*S*-phenylglycine *t*-butylester), an inhibitor of γ-secretase which prevents cleavage of Notch1 at the S3 site to release the ICD. Acute treatment with DAPT results in focal adherens junctions and disorganized cortical actin fibers (Fig. 3G-I; Supplemental Fig. 5C, D). Further, visualizing live actin dynamics following treatment with DAPT revealed dissolution of cortical actin fibers within 30 minutes (Supplemental Fig. 5E), a rapid response that further supports a transcription-independent Notch1 function. Treatment of *NOTCH1^KO^* monolayers with Erlotinib did not prevent focal adherens junctions, confirming that the adherens junction phenotype is independent of EGFR kinase activity (Supplementary Figure 5F). Altogether, these results identify that Notch1 cortical signaling functions to stabilize epithelial adherens junctions and cortical actin organization.

### Localization and cleavage of Notch1 at lateral cell-cell contacts

Phenotypes in *NOTCH1^KO^* ducts present during lumen expansion (Figure 1) and *NOTCH1^KO^* cells have impaired columnar morphology (Figure 2), indicating that Notch1 cortical signaling may play a role in stabilizing adherens junctions as cells reach confluence and undergo apical-basal restructuring. To further understand the mechanistic contribution of Notch1 cortical signaling to this morphodynamic change, we assessed the localization of endogenous Notch1 within mammary epithelia cultured on compliant ECM hydrogels at distinct morphogenic timepoints ranging from low confluence (LC), where cells are surrounded by other cells yet remained elongated, to high confluence (HC) where cells are surrounded and cuboidal but still flat, to polarized (P) where cells had adopted a columnar morphology. During this transition, immunofluorescence staining revealed Notch1 progressively accumulates at cell-cell interfaces (Figure 4A, B). High magnification confocal micrographs further showed that E-cadherin most strongly localizes to apical domains in the polarized state, while Notch1 and cortical actin intensity is highest at lateral cell membranes (Figure 4C). This lateral localization is consistent with a recent study reporting that Notch1 activity is limited by the formation of lateral membrane contacts and adherens junctions during cellularization of the embryonic *Drosophila* syncytium (Falo-Sanjuan and Bray, 2021).

**Figure 4.**
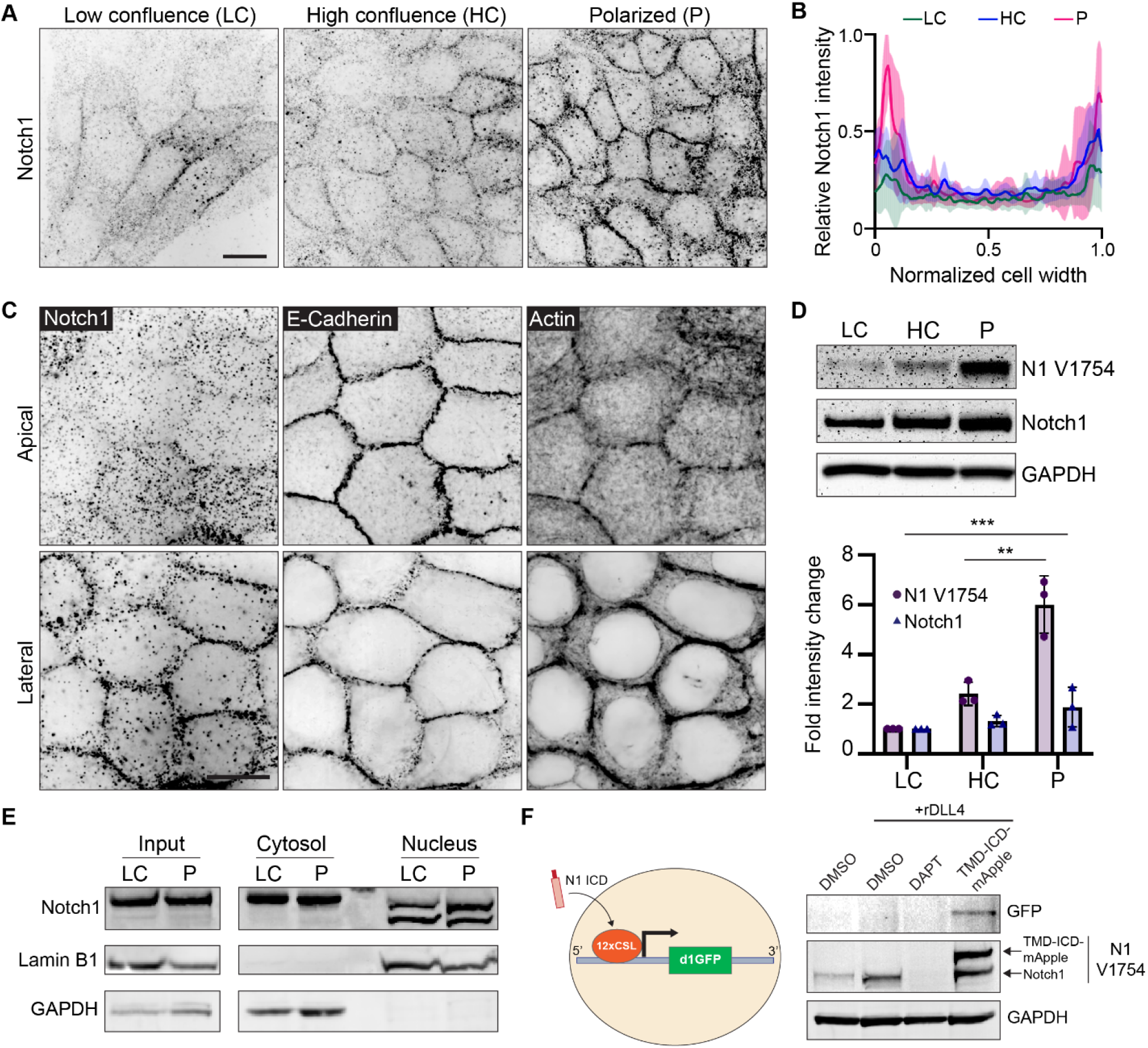
Localization and proteolytic activation of Notch1 at lateral cell-cell contacts. (A) Immunofluorescence micrographs of Notch1 (black) in wild type MCF10A in low confluence (LC), high confluence (HC), and polarized (P) states. Scale bar, 20 μm. (B) Quantification of relative Notch1 intensity across the width of the cell. n = 8 cells from three independent experiments. (C) Representative SoRa immunofluorescence micrographs of wild type monolayers immunostained for E-Cadherin and Notch1 and labeled with phalloidin. Top row: apical domain. Bottom row: lateral domain. Scale bar, 20 μm. (D) Top: Western blot of wild type lysates from the indicated monolayer states, immunoblotted for cleaved Notch1 V1754 (N1 V1754), total Notch1, and GAPDH. Bottom: Quantification of fold change in N1 V1754 and total Notch1 band intensities. n = 3 independent experiments. (E) Western blot of cytosolic and nuclear fractions from wild type monolayer lysates in low confluence (LC) and polarized (P) states, immunoblotted for total Notch1, Lamin B1, and GAPDH (F) Left: Schematic of Notch1 transcriptional destabilized GFP reporter (d1GFP). Right: Western blot of lysates from wild type monolayers treated with DMSO, DMSO + rDLL4, DAPT + rDLL4, or overexpressing constitutively active form of Notch1 (TMD-ICD-mApple), immunoblotted for GFP, N1 V1754, and GAPDH. Western blots are representative of three independent experiments. For plot in D, mean ± SEM; one-way ANOVA with Tukey’s post-hoc test, **p<0.01, ***p<0.001.

We next examined whether Notch1 junctional accumulation correlated with functional changes in ICD S3 cleavage or transcriptional activity. Western blot analysis of monolayers transitioning between low confluence and polarized states revealed that Notch1 junctional accumulation is coincident with a six-fold increase in γ-secretase-mediated cleavage of ICD (cleavage-specific Notch1 V1754 antibody) with only a marginal increase in total Notch1 protein levels (Figure 4D). Despite substantial increases in cleaved ICD, the amount of ICD within isolated nuclear fractions does not markedly change between low confluence and polarized states (Figure 4E). Interestingly, ICD within nuclear fractions presents as two lower molecular weight bands relative to ICD in cytosolic fractions, which is consistent with intracellular post-translational regulatory mechanisms directing ICD function after cleavage (Antfolk et al., 2019).

To further investigate if increased ICD cleavage in the polarized state leads to ICD-dependent transcription, we generated mammary epithelial cells stably expressing a fluorescent proteinbased Notch transcription reporter consisting of twelve CSL-binding motifs coupled to a destabilized GFP (d1GFP) that has an approximate half-life of 1-2 hours (Hansson et al., 2006). Despite increased levels of cleaved ICD, polarized monolayers have no detectable GFP expression. Coating hydrogels with recombinant Notch ligand Delta Like Canonical Notch Ligand 4 (rDll4) increases ICD cleavage, but similarly lacks reporter GFP expression. However, expressing a constitutively active form of Notch1 lacking the ECD (TMD-ICD-mApple) (Chiang et al., 2006; Polacheck et al., 2017) at levels approximately two-fold endogenous Notch1 is sufficient to stimulate reporter GFP expression (Figure 4F). Altogether, these data indicate that as cells reach confluence and initiate apical-basal restructuring, Notch1 is localizes to lateral cell-cell contacts and the ICD is proteolytically cleaved. This increase in cleaved ICD does not lead to higher levels of ICD in nuclear fractions or the expression of a Notch transcriptional reporter, which can be engaged by expression of a constitutively active form of Notch1. Along with truncation of the ICD and expression of dnMAML, this observed elevated proteolytic activation absent a robust transcriptional response further affirms an important role for Notch1 cortical signaling in stabilizing adherens junctions in epithelial monolayers.

### Notch1 cortical signaling functions through FAM83H to stabilize adherens junctions and regulate duct architecture

To identify molecular pathways associated with the localization and/or adherens junction stabilizing function of Notch1 cortical signaling at lateral cell contacts, we unbiasedly profiled differential Notch1 interacting proteins within low confluence and polarized monolayers using Notch1 immunoprecipitation, SDS-PAGE and Coomassie staining, and mass spectrometry (Figure 5A, Supplementary Figure 6A). Notably, while this approach identified several distinct interactions, Notch transcriptional effectors MAML1/2 and RBPJ were not identified, supporting the observed lack of increase in nuclear ICD within polarized monolayers (Figure 4E). One prominent band (~150 kDa) isolated from polarized monolayers was identified as FAM83H from the FAM83 family of oncogenes (Snijders et al., 2017). Co-immunoprecipitation and western blot confirmed a FAM83H-Notch1 interaction that increases three-fold as monolayers progress from low confluence to polarized states (Figure 5B).

**Figure 5.**
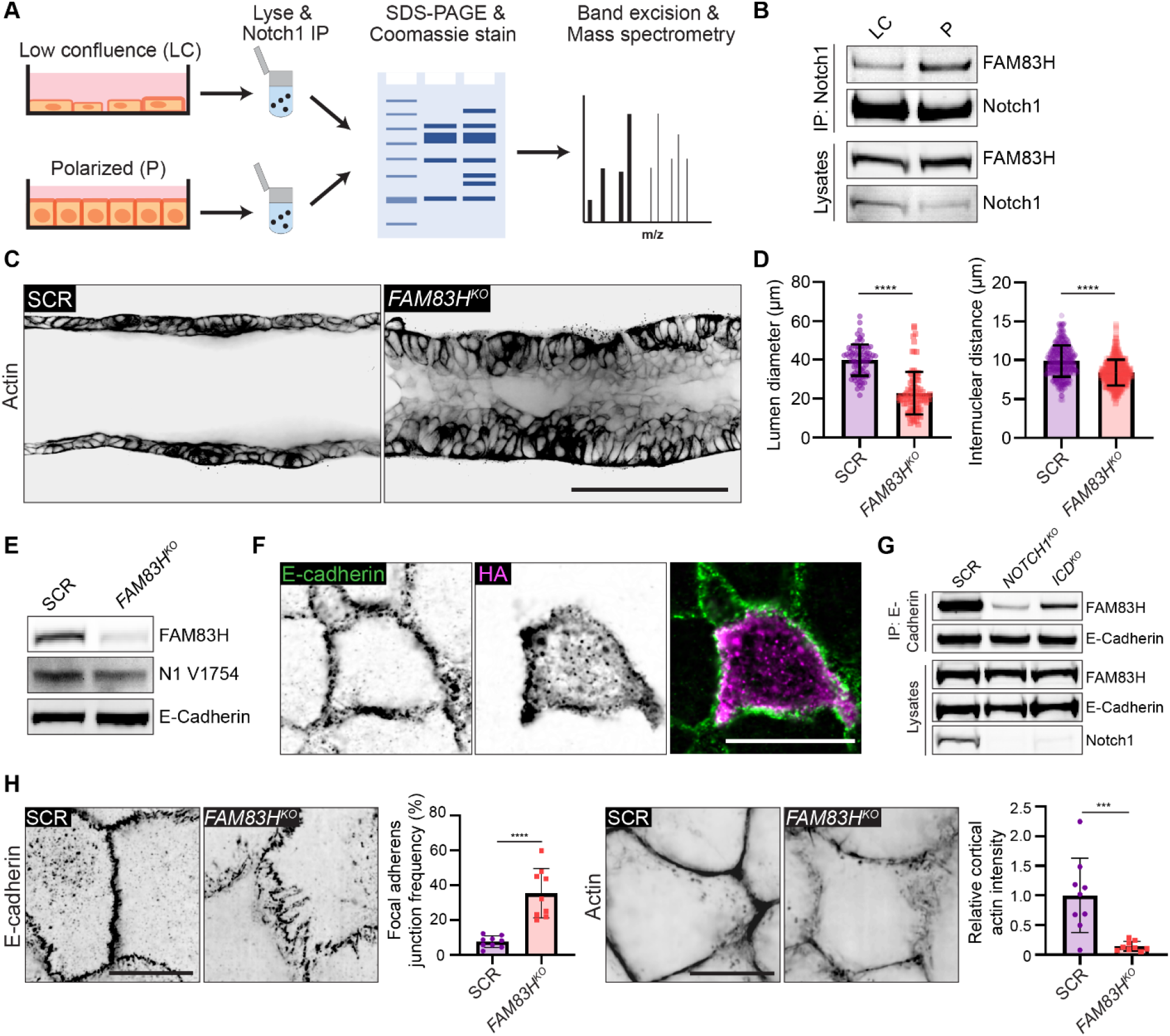
Notch1 cortical signaling functions through FAM83H to stabilize adherens junctions and control duct architecture. (A) Schematic of mass spectrometry workflow to identify monolayer state-dependent, differential Notch1 protein-protein interactions. (B) Western blot of immunoprecipitated Notch1 from low confluence (LC) or polarized (P) monolayers immunoblotted for FAM83H and Notch1. (C) Representative medial confocal slice fluorescence micrographs of SCR and *FAM83H^KO^* ducts labeled with phalloidin (black). Scale bar, 100 μm. (D) Left: Quantification of duct lumen diameter. n ≥ 9 average lumen diameters from three independent experiments. Right: Quantification of internuclear distances measured from Hoechst labeled ducts. n ≥ 50 internuclear distances per duct from three independent experiments. (E) Western blot of lysates from SCR or *FAM83H^KO^* monolayers immunoblotted for FAM83H, cleaved Notch1 V1754 (N1 V1754), and E-Cadherin. (F) Immunofluorescence micrographs of HA-FAM83H expressing cell immunostained for E-Cadherin (green) and HA (magenta). Scale bar, 20 μm. (G) Western blot of immunoprecipitation of E-Cadherin from SCR, *NOTCH1^KO^*, and *ICD^KO^* monolayer immunoblotted for FAM83H and E-Cadherin. (H) Left: Immunofluorescence micrographs of SCR and *FAM83H^KO^* cells immunostained with E-Cadherin (black) and the corresponding quantification of the frequency of focal adherens junctions. Scale bar, 10 μm. n ≥ 9 fields of view from three independent experiments. Right: Fluorescence micrographs of SCR and *FAM83H^KO^* monolayers labeled with phalloidin (black) and the corresponding quantification of cell-cell junction cortical actin intensity. Scale bar, 10 μm. n ≥ 9 fields of view from three independent experiments. Western blots are representative of three independent experiments. For all plots, mean ± SEM; two-tailed unpaired t test, ***p<0.001, ****p<0.0001.

The cellular function of FAM83H is not well understood, despite reported roles in development, cancer, and intermediate filament dynamics (Kim et al., 2008; Kim et al., 2019; Kuga et al., 2016). We first investigated whether FAM83H contributes to the morphogenic phenotypes associated with loss of Notch1 cortical signaling by generating 3D ducts from mammary epithelial cells depleted of FAM83H by CRISPR-Cas9 (*FAM83H^KO^). FAM83H^KO^* phenocopies key morphologic signatures associated with *NOTCH1^KO^* ducts, namely a tortuous duct architecture that is driven by lumen cell in-filling, multilayering, and increased cell packing (Figure 5C, D; Supplementary Figure 6B). *FAM83H^KO^* does not abolish Notch1 ICD cleavage, Notch1 localization to cell-cell contacts, or E-cadherin expression levels (Figure 5E) suggesting FAM83H functions as a downstream arm of the Notch1 cortical pathway. We were unable to identify a commercially available antibody to visualize endogenous FAM83H localization, thus we isolated the FAM83H coding sequence and generated an HA-tagged FAM83H expression construct. HA-FAM83H localizes to cell-cell contacts and cytoplasmic aggregates in cells within polarized monolayers (Figure 5F). Consistent with identification in the E-cadherin adhesome (Guo et al., 2014) and a function downstream of Notch1 cortical signaling, FAM83H co-immunoprecipitates with E-cadherin but this interaction is lost in *NOTCH1^KO^* and maintained in *ICD^KO^* cells (Figure 5G). Furthermore, *FAM83H^KO^* results in focal adherens junctions and disordered cortical actin, resembling phenotypes observed upon loss of Notch1 cortical signaling in *NOTCH1^KO^* (Figure 5H; Supplemental Figure 6C). Together, these data provide evidence for a novel interaction between Notch1 and FAM83H and identify FAM83H as a new Notch1 cortical signaling effector that functions in the stabilization of epithelial adherens junctions and cortical actin organization.

## DISCUSSION

Notch signaling broadly controls developmental and homeostatic morphogenic processes that involve coordinated changes in cell architecture and gene expression; however, how a singular canonical transcriptional pathway is able to produce such diverse biological output is unclear (Bray, 2016). The possibility of direct links from Notch to cell-cell adhesion and the actin cytoskeleton has been suggested in past studies on axon guidance, keratinocyte motility, *Drosophila* embryogenesis, sprouting angiogenesis (Crowner et al., 2003; Lowell and Watt, 2001; Major and Irvine, 2005; Zakirov et al., 2021), and our work describing a transcription-independent Notch1 cortical signaling pathway that regulates adherens junction assembly and barrier function in vascular endothelial cells in response to shear stress (Polacheck et al., 2017). Our study here reveals a new context by which Notch1 controls the tissue architecture of an organotypic human mammary ductal epithelium through a mechanism independent of the Notch1 ICD and transcriptional signaling.

We report that Notch1 controls epithelial cell architecture and proliferation by regulating adherens junctions independent of transcription. This is supported by genetic perturbation that specifically truncates endogenous Notch1 ICD and a dominant negative approach to globally suppress Notch transcription, which mitigates potential changes in transcriptional signaling from other Notch receptors that might occur between *NOTCH1^KO^ and ICD^KO^*. Cell proliferation increase occurs noncell autonomously through enhanced sensitivity to EGF and internalization of EGFR. This is consistent with Notch1 tumor suppressor function in murine skin, where *Notch1^-/-^* epidermal keratinocytes promote tumorigenesis non-cell autonomously by impairing skin barrier integrity which drives dysplasia through the generation of a mitogenic, wound-like microenvironment (Demehri et al., 2009; Nicolas et al., 2003). While offering important insight into an unappreciated arm of Notch1 signaling, this study also illuminates a new mechanism by which cell-cell adhesion may be dynamically regulated. There is considerable evidence *in vivo* and *in vitro* that modulation of Notch1 activation converges on adherens junction dynamics (Bentley et al., 2014; Falo-Sanjuan and Bray, 2021; Grammont, 2007; Polacheck et al., 2017), and we are just beginning to understand the molecular underpinnings. Our previous work identified the Notch1 TMD as the critical domain mediating protein-protein interactions which underlie adherens junction alterations in the vascular endothelium (Polacheck et al., 2017), and our results here are consistent with a central function for Notch1 TMD. Recent studies have shown that transmembrane protein–protein interactions are critical determinants of subcellular distribution, clustering, and signaling (Coon et al., 2015; Diaz-Rohrer et al., 2014; Lewis et al., 2019). This work advances our understanding of transcription-independent Notch1 function by identifying a previously unappreciated interaction with FAM83H, whereby FAM83H recruitment to E-cadherin is dependent on Notch1 cortical signaling. We find that HA-FAM83H localizes to cell-cell contacts and intracellular aggregates in polarized epithelial monolayers. This is in agreement with reports describing FAM83H localization to cell-cell interfaces *in vivo* (Kuga et al., 2016) and posit FAM83H functions as a peripheral membrane protein (Ding et al., 2009). Still, the nature of FAM83H interaction and recruitment within the context of Notch1 cortical signaling is yet to be determined.

Notch signaling depends on the size and geometry of the contact sites in sender-receiver cell models (Shaya et al., 2017). Here, within a homogenous epithelial monolayer, we report Notch1 cortical signaling is uniformly engaged as Notch1 accumulates at lateral cell-cell interfaces and is proteolytically cleaved during epithelial apical-basal columnar restructuring. Notch1 accumulation at lateral interfaces independent of protein level change suggests active recruitment through mechanisms linked to this change in cellular architecture, and is consistent with a recent report of Notch regulation by the polarity protein Par3 *in vivo* (Wu et al., 2022). Moreover, we observe a substantial increase in γ-secretase cleaved ICD during this transition, yet this increase in cleaved ICD does not lead to more nuclear-localized ICD or transcription, suggesting intricate mechanisms gate transcriptional activity of cleaved ICD. While the distinct molecular weights observed in nuclear biochemical fractions indicate post-translational ICD regulation, another potential mechanism is an additional requirement for nuclear mechanotransduction. In this model, forces on the nucleus gate ICD transcription, either through regulation of nuclear pore transport or altered chromatin accessibility, as was proposed for Notch during mesoderm invagination in *Drosophila* gastrulation (Falo-Sanjuan and Bray, 2022).

How FAM83H contributes to adherens junction stability is an outstanding question. One of the few prescribed functions of FAM83H is the regulation of keratin intermediate filament dynamics through casein kinase I (Kim et al., 2008; Kim et al., 2019; Tokuchi et al., 2021). A recent report described a role for keratin-desmosome networks in organizing the cortical actin cytoskeleton to control epithelial cohesion and limit tensile stress on adherens junctions (Prechova et al., 2022), which aligns with adherens junction phenotypes upon loss of Notch1 cortical signaling. Interestingly, loss of function mutations in *Fam83h* and *Notch1* both cause amelogenesis imperfecta and a failure to form desmosomes between ameloblast and stratum intermedium layers in the mouse incisor (Jheon et al., 2016; Kim et al., 2008). Similarly, *Fam83h^-/-^* mice die by postnatal day 14 and have visible skin defects (Wang et al., 2016). *Notch1^-/-^* keratinocytes impair epidermal barrier integrity, which is a tissue property that is regulated by desmosome-keratin networks (Johnson et al., 2014). Keratin filaments interact with Notch1 to regulate colonic epithelial proliferation and differentiation through an unknown mechanism (Lahdeniemi et al., 2017). It is therefore plausible that reciprocal interactions between Notch1 and desmosome-keratin networks, which involve FAM83H, may contribute to the maintenance of epithelial cell-cell adhesion and differentiation.

Together, our work offers new insights into Notch1 signaling and regulation, how cell-cell adhesions are dynamically regulated, and a model in which transcriptional and adhesive programs might be coordinated by a single receptor. The convergence of transcription-independent Notch1 cortical signaling on cell-cell adhesion regulation may explain skin barrier defects associated with the tumor suppressive function of Notch1, as well as tissue jamming/fluidization events occurring during developmental morphogenesis. Identifying ways to isolate the Notch1 cortical pathway from the transcriptional pathway may therefore provide new opportunities to instruct development and to treat associated complications.

## Acknowledgements

The authors thank Young-wook Jun, Lakyn Mayo, Diane Barber, Torsten Wittmann, and Yi Liu for critical assistance and feedback on the manuscript. This work was supported by NIH grants R00CA226366 and R21AG072232 and the UCSF Program for Breakthrough Biomedical Research. The Nikon CSU-W1 SoRa microscope was obtained through NIH shared equipment grant S10OD028611-01.

## Author Contributions

MLK conceived the study. MJW and MLK performed the experiments with assistance from KJ and TS. MJW and MLK wrote the manuscript with feedback from all authors.

## METHODS

### Cell culture

Human mammary epithelial cells (MCF10A, ATCC) were maintained in growth medium consisting of DMEM/F12 (Gibco), 5% horse serum (Invitrogen), 20 ng/ml EGF (PeproTech), 0.5 mg/ml hydrocortisone (Sigma), 100 ng/ml cholera toxin (Sigma), 100 U/ml penicillin, 100 μg/ml streptomycin (Invitrogen) and 10 μg/ml insulin (Sigma). Human bronchial epithelial cells (16HBE14o-, Sigma) were maintained in medium consisting of α-MEM (Sigma), 10% fetal bovine serum (Peak), 100 U/ml penicillin, 100 μg/ml streptomycin (Invitrogen) and 2 mM L-glutamine (Sigma). Human colorectal adenocarcinoma cells (Caco-2, ATCC) were maintained in medium consisting of DMEM (Sigma), 100 U/ml penicillin, 100 μg/ml streptomycin (Invitrogen) and 20% fetal bovine serum (Peak). Human HEK-293T cells (Clonetech) were maintained in DMEM (Sigma), 10% fetal bovine serum (Peak), 2 mM L-glutamine, 100 U/ml penicillin, 100 μg/ml streptomycin (Invitrogen), and 1 mM sodium pyruvate (Gibco). MCF10A, 16HBE14o-, and Caco-2 cells were used at passages 2-12 and all cell types were maintained at 37°C in 5% CO2 in a humidified incubator. Cell numbers were counted at passage using the Countess 3 automated cell counter (Invitrogen). Cell-line authentication (performance, differentiation, and STR profiling) was provided by ATCC, Sigma, and Clonetech. All cells were routinely tested for *Mycoplasma* via PCR test (Applied Biological Materials).

### Antibodies and reagents

Antibodies against Notch1 ICD (D1E11, 1:100 IF, 1:1000 WB), Notch1 V1744 (D3B8, 1:500 WB), EGFR (D38B1, 1:100 IF, 1:1000 WB), pEGFR (D7A5, 1:1000 WB), GFP (D5.1, 1:1000 WB), non-phospho (Active) β-Catenin (D13A1, 1:1000 WB), GAPDH (14C10, 1:10000 WB) and YAP (D8H1X, 1:200 IF) were from Cell Signaling Technologies. β-catenin antibody (14, 1:1000 WB) was from BD Biosciences. E-cadherin antibody (HECD-1, 1:1000 IF, 1:1000 WB) was from Takara Bio. Notch1 ECD (ABS90, 1:1000) was from Millipore. FAM83H antibody (1:1000 WB) was from Bethyl Laboratories. Lamin B1 antibody (12987-1-AP, 1:1000) was from Proteintech. Rhodamine phalloidin and Alexa Fluor 488, 568 and 647 goat anti-mouse and anti-rabbit IgG secondary antibodies (1:400) were from Invitrogen. Alexa Fluor 647 azide and EdU cell proliferation kit was from Invitrogen. Hoescht and DAPT were from Sigma.

### Lentiviral-mediated CRISPR-Cas9 editing

Stable CRISPR-modified primary MCF10A, 16HBE14o-, and Caco-2 cell lines were generated using the lentiCRISPRv2 system using our previously established protocols (Kutys et al., 2020; Polacheck et al., 2017). Specific guide RNAs were cloned into the BsmBI site of plentiCRISPRv2: SCR, GTATTACTGATATTGGTGGG; *NOTCH1^KO^*, CGTCAGCGTGAGCAGGTCGC; *ICD^KO^*, TGCTGTCCCGCAAGCGCCGG; *FAM83H^KO^*, GGACAACCCACTGGCACCC. sgRNA-containing plentiCRISPRv2 plasmids were co-transfected with psPAX2 (Addgene plasmid #12260) and pMD2.G (Addgene plasmid #12259) packaging plasmids into HEK-293T cells using calcium phosphate transfection. After 48 hours, viral supernatants were collected from the culture dish, concentrated using 4x lentivirus concentrator (PEG-IT), and resuspended in PBS. MCF10A, 16HBE14o-, and Caco-2 cells were transduced in their corresponding growth medium overnight and given fresh medium the following morning. At 48 hours post transduction, cells were passaged and plated in 6-well plates at 1.25 x 10^5^ cells per well and selected with 2 μg/ml puromycin for 4 days. All CRISPR modifications were verified by western blot.

### Microfluidic device design and fabrication

The microfluidic device contains four main ports (two basal ports and two luminal ports) along with a central hydrogel-containing compartment. The basal ports functioned as media reservoirs containing all supplemental components of MCF10A culture medium and luminal ports functioned as inlet and outlet reservoirs to maintain lumen pressure. Between the hydrogel compartment and the basal ports are a row of PDMS pillars which span the height of the hydrogel compartment and function to contain the hydrogel within that central region. The silicon master was fabricated using photolithography methods previously described (Polacheck et al., 2019) however UV exposure steps were performed on an Alvéole PRIMO micropatterning system. Individual microfluidic devices were generated using soft lithography. Polydimethylsiloxane (PDMS, Sylgard 184, Dow-Corning) was mixed at a ratio of 10:1 (base:curing agent) and cured at 60 °C on a silicon master. The PDMS was cut from the silicon master, trimmed, and surface-activated by plasma treatment for 30 seconds at 300 mTorr. Devices were then bonded to glass coverslips and surface treated with 0.01% poly-L-lysine for two hours and 1% glutaraldehyde for 15 minutes. Devices were then washed 3x with water and sterilized in 70% ethanol for 30 minutes. Steel acupuncture needles (160 μm diameter, Tai-Chi) were inserted into each device and devices were placed into a vacuum desiccator for 60 minutes. Unpolymerized ECM mixture containing 70% neutralized collagen type I (Corning) and 30% growth factor reduced Matrigel (Corning) was injected into the hydrogel compartment via the extracellular matrix-loading port. Collagen type I solution was buffered with 10x DMEM, 10X reconstitution buffer containing 0.2 M HEPES and 0.26 M Sodium bicarbonate, titrated to a pH of 7.6 with NaOH, and brought to a final concentration of 2.5 mg/ml collagen I in PBS. Once the unpolymerized hydrogel was injected into the central compartment, the devices were incubated at 37 °C for 25 minutes. Following polymerization of the hydrogel, all ports were filled with sterile PBS and devices were left overnight at 37 °C. Steel acupuncture needles were removed from the devices to create 160 μm-diameter channels in the hydrogel and the outer edges of the PDMS were sealed with vacuum grease (Dow-Corning). Devices were kept hydrated with sterile PBS and maintained at 37 °C in 5% CO_2_ humidified air.

### Organotypic mammary duct fabrication and culture

MCF10A were harvested with 0.05% Trypsin-EDTA (Gibco) and centrifuged at 200 *g* for 3 minutes. Cells were resuspended at 2 x 10^6^ cells/ml in assay medium containing DMEM/F12 (1:1, Gibco) supplemented with 5% horse serum (Invitrogen), 5 ng/ml EGF (PeproTech), 0.5 μg/ml hydrocortisone (Sigma), 100 ng/ml cholera toxin (Sigma), 100 U/ml penicillin, 100 μg/ml streptomycin (Invitrogen) and 10 μg/ml insulin (Sigma). Following resuspension, 70 μl of cell suspension was added to the two luminal ports to allow for perfusion and adherence of cells to the channel surface. Cells were perfused with occasional device flipping to coat the top of the channel for approximately 15 minutes or until 70% cell coverage prior to exchanging with growth medium. Luminal and basal ports were then filled with fresh assay medium, and devices were incubated at 37 °C in 5% CO2 humidified air. Assay medium was replenished in all ports for days 1 and 2 post-seeding. After a confluent tissue formed (typically day 3), assay medium was introduced to basal ports and lumen ports were changed to a base medium daily containing only DMEM/F12 (Invitrogen) supplemented with 100 U/ml penicillin and 100 μg/ml streptomycin (Invitrogen). Dead cells accumulating in the lumen were removed by perfusion and medium changes were performed every 24h.

### Immunofluorescence

Duct tissues were fixed in 4% paraformaldehyde in PBS supplemented with calcium and magnesium (PBS++) for 15 minutes on a rocker at 37°C, rinsed three times with PBS, and permeabilized in 0.1% Triton X-100 for 1 hour on a rocker. Ducts were rinsed three times with PBS and blocked with 2% BSA in PBS overnight on a rocker at 4°C. Primary and secondary antibodies were added to all device ports in 2% BSA in PBS and devices were placed on a rocker overnight at 4°C. Ducts were rinsed three times over 3 hours with PBS between primary and secondary antibody treatments.

For immunofluorescence of 2D monolayers on hydrogels, cells (MCF10A, 16HBE14o-, and Caco-2) were plated on 18 mm glass coverslips coated with an 80:20 collagen type I:Matrigel hydrogel. Coverslips were surface activated by plasma treatment for 30 seconds, coated with 0.01% poly-L-lysine for 45 minutes, and washed three times with PBS. Collagen type I solution and Matrigel mixture was prepared as described in device fabrication methods. Once prepared, 40 μL of hydrogel was added and spread around the surface of each coverslip and incubated at 37 °C for 20 minutes. Coverslips with polymerized hydrogel were added to 12-well plates and medium was added to each well. All epithelial cell types were resuspended at 2 x 10^6^ cells/ml in their corresponding culture medium and 1.25 x 10^5^ cells were added to each well. Once cells reached a confluent monolayer, coverslips were fixed in 4% paraformaldehyde in PBS++ for 15 minutes at 37 °C, rinsed three times with PBS, and permeabilized in 0.1% Triton X-100 for 30 minutes. Coverslips were washed three times with PBS and blocked in 2% BSA in PBS for 1 hour. Primary and secondary antibodies were applied in 2% BSA in PBS for 1-2 hours at room temperature or overnight at 4 °C and rinsed three times over 30 minutes with PBS between each treatment. For immunofluorescence imaging of mammary epithelial ducts, images were acquired on either a Yokogawa CSU-X1 spinning disk confocal on a Nikon Ti-E microscope or a Yokogawa CSU-W1/SoRa spinning disk confocal on a Ti2 inverted microscope stand (Nikon) with a 40x 1.25 water immersion lens (Nikon). For live-cell or immunofluorescence imaging of epithelial monolayers, images were acquired on the Yokogawa CSU-W1/SoRa spinning disk confocal system in SoRa mode with a 60x 1.49 NA oil immersion lens (Nikon). Fluorescence images were adjusted for contrast and brightness using ImageJ.

### Live cell imaging

Confluent monolayers were labeled for two hours with SPY650-FastAct (Cytoskeleton Inc) by adding 1x of the probe to growth medium. Immediately prior to imaging, DAPT (10 μM) was added to the culture medium. For the epithelial ducts, 1x SPY650-FastAct (Cytoskeleton Inc) was applied for four hours prior to imaging. All images were acquired on a Yokogawa CSU-X1 spinning disk confocal on a Nikon Ti-E microscope equipped with an imaging chamber equilibrated to 37 °C in 5% CO_2_ humidified air.

### Image processing and analysis

To quantify average lumen size, confocal micrographs of a midline slice of duct tissues stained with phalloidin were processed in ImageJ and the width of the lumen was measured at 10 distinct locations along the duct. Epithelial duct internuclear distances were determined by measuring the distance from the midpoint of one nucleus to the midpoint of its neighboring nuclei in ImageJ. Variance in diameter was quantified by binarizing phase images of ducts through global intensity thresholding in MATLAB. Outer boundaries of each duct were determined by marking the first and last white pixel in each column of the image matrix, and the diameter along the duct was compared to the mean diameter. To quantify the number of EdU positive nuclei, the fractional percentage (% positive nuclei) was calculated by dividing the number of EdU positive nuclei by the total number of nuclei. Regions of multilayering were defined by the number of regions within a field of view that had three or more multilayered nuclei above the monolayer plane. To quantify the number of focal adherens junctions, confocal micrographs of epithelial monolayers immunostained for E-Cadherin were processed in ImageJ and the total number of cell-cell junctions was measured. Focal adherens junctions were defined as cell-cell junctions that had a discontinuous, jagged, and non-linear phenotype and the fractional percentage was calculated by dividing the number of focal adherens junctions by the total number of junctions. To quantify relative cortical actin intensity, line profiles were drawn through the short axis of the cell, passing through the nucleus. The intensity of phalloidin-labelled cells was plotted along lines. Cortical actin was defined as the area under the peak at cell–cell junctions normalized to total area under the curve. Cell height was determined by measuring the distance in ImageJ from the basal surface to the apical surface in orthogonal projection fluorescence micrographs of monolayers stained with phalloidin. To quantify Notch1 localization, line profiles were drawn through the short axis of the cell, passing through the nucleus. Notch1 signal intensity was plotted along this line and intensity values and length were normalized to their respective maxima. To quantify EGFR localization, fluorescence micrographs of monolayers immunostained for EGFR were segmented into individual cells and thresholded by Otsu’s method in MATLAB. Cytoplasmic to junctional EGFR ratio was defined as the total number of pixels within the cell interior divided by those at cell-cell interfaces.

### Cloning and qPCR

Cells were lysed with cold TRI reagent (Sigma) following the prescribed protocol. RNA extraction was performed according to the manufacturer’s protocol (Zymogen Direct-zol RNA MiniPrep Kit). cDNA libraries were generated with SuperScript II (FisherFull length human FAM83H cDNA was amplified from the library by PCR using primers containing overhangs to allow for subsequent Gibson assembly of HA-tagged FAM83H in a modified pRRL lentiviral expression vector. FAM83H cDNA sequence was verified by whole plasmid sequencing. Realtime PCR was performed in 20 μl reactions using the SYBR Green Master Mix protocol (ThermoFisher) and BioRad thermocycler. The following qPCR primers were obtained from IDT: hPSMB2_F ACTATGTTCTTGTCGCCTCCG, hPSMB2_R CTGTACAGTGTCTCCAGCCTC, hHES1_F CCAAGTGTGCTGGGGAAGTA, hHES1_R CACCTCGGTATTAACGCCCT, hHEY1_F CTGAGCAAAGCGTTGACA, and hHEY1_R TCCACCAACACTCCAAA. Relative gene expression levels were calculated by -ΔΔCT in which ΔCT is the difference between the CT value of the gene of interest (HES1 or HEY1) and the CT value of the housekeeping gene (PSMB2) and ΔΔCT is the difference between the experimental condition and the control.

### Immunoblotting

For western blot of monolayers transitioning between low confluence and polarized states, cells were plated into individual wells of a six well plate in assay medium at 1.25 x 10^5^ cells/well. Assay media was changed daily. Cells were lysed in buffer described below at 24 hours for low confluence, 48 hours for high confluence, and 72 hours for polarized monolayer states. Monolayers cultured in assay medium (unless otherwise noted) were rinsed with PBS and lysed with 50 mM Tris-HCl, pH 7.4, 150 mM NaCl, 1% Triton X-100, 0.1% SDS, 0.5% Sodium deoxycholate and 1.5x protease and cOmplete Protease Inhibitor (ThermoFisher). Lysates were passed through 21G syringe 10 times and incubated on ice for 10 minutes prior to centrifugation at 4°C for 10 minutes at 13000x*g*. Lysate protein content was normalized using a BCA protein assay kit (Prometheus) and samples were denatured with 1x NuPAGE LDS Sample Buffer (Life Technologies) containing 5%β-Mercaptoethanol. Denatured lysates were analyzed by SDS-PAGE and gels were transferred to PVDF membranes using a Mini Trans-Blot Cell (Bio-Rad). Membranes were blocked in 5% non-fat dry milk in TBS containing 0.1% Tween-20 (TBST) for one hour at room temperature. Primary antibodies were applied to membranes in blocking buffer overnight at 4°C. Membranes were washed three times over 30 minutes with TBST. IRDye donkey anti-rabbit and anti-mouse IgG secondary antibodies (1:10000) (LI-COR) were incubated in blocking buffer for two hours at room temperature. Membranes were washed three times over 30 minutes with TBST. All immunoblots were imaged using an Odyssey CLx LI-COR Imaging System and quantified using ImageJ. Immunoblots were adjusted for brightness and contrast using ImageJ and (unless otherwise noted) intensity values were normalized to the GAPDH loading control. All uncropped western blots are provided in Supplementary Figure 7.

### Traction force microscopy

Polyacrylamide gels of desired stiffness were made by adjusting acrylamide and bisacrylamide stock solution (Bio-Rad Laboratories, Hercules, CA) concentrations (Chopra et al., 2018). A solution of 40% acrylamide, 2% bisacrylamide and 1xPBS was polymerized by adding tetramethylethylene diamine (Fisher BioReagents) and 1% ammonium persulfate. A droplet of the gel solution supplemented with 0.2 μm fluorescent beads solution (Molecular Probe, Fisher Scientific) was deposited on a quartz slide (Fisher Scientific) and covered with a 25-mm glass (Fisher) coverslip pretreated with 3-aminopropyltrimethoxysilane (Sigma-Aldrich) and glutaraldehyde (Sigma-Aldrich). After polymerization, the gel surface attached to the quartz slide was functionalized with fibronectin via EDC-NHS chemistry. Briefly, the gel surface was activated in a UV-Ozone cleaner (Jelight) for 2 minutes, detached from the quartz slide, soaked in a solution with EDC and NHS for 15 minutes and incubated with 50 μg/ml fibronectin solution at 37°C for 2 hours. The gel was sterilized and stored in 1X PBS before cell seeding. The traction forces exerted by epithelial monolayers on the polyacrylamide gel substrates were computed by measuring the displacement of fluorescent beads embedded within the gel. Briefly, images of bead motion near the substrate surface, distributed in and around the contact region of a single cell (before and after cell detachment with 10% sodium dodecyl sulfate, were acquired with Yokogawa CSU-21/Zeiss Axiovert 200M inverted spinning disk microscope with a Zeiss LD C-Apochromat 40×, 1.1 N.A water-immersion objective and an Evolve EMCCD camera (Photometrics). The traction stress vector fields were generated using an open source package of FIJI plugins (Tseng et al., 2012).

### Nuclear/cytosolic fractionation

Monolayers cultured in 10 cm plates in growth medium were rinsed once with PBS++ and then scraped into cold PBS++. Cells were pelleted by centrifugation at 500xg and supernatant was aspirated. Cells were resuspended and swollen using ice cold hypotonic lysis buffer (20mM Tris-HCl pH 7.4, 10mM KCl, 2mM MgCl2, 1mM EGTA, 0.5mM DTT, 1.5x cOmplete Protease Inhibitor). Cells were then lysed by adding Triton X-100 to 0.1% and incubated on ice for three minutes. Lysate was centrifuged at 1000 g to pellet nuclei and supernatant (cytoplasmic fraction) was collected. Supernatant was clarified by centrifugation at 15,000 g for 3 minutes. Nuclear pellet was resuspended in cold isotonic buffer (20mM Tris-HCl pH 7.4, 150mM KCl, 2mM MgCl2, 1mM EGTA, 0.3% Triton X-100, 0.5mM DTT, cOmplete Protease Inhibitor) and incubated for 7 minutes. Nuclei were centrifuged at 1000 g for three minutes and supernatant was aspirated. Nuclei were resuspended in cold RIPA buffer (50mM Tris-HCl pH 7.4, 150mM NaCl, 0.1% SDS, 0.5% sodium deoxycholate, 1% Triton X-100, cOmplete Protease inhibitor) and were incubated on ice for 30 minutes. Nuclei were then centrifuged at 2000 g for 3 minutes and supernatant (nucleosol fraction) was collected. Lysate protein content was normalized for each fraction using a BCA protein assay kit (Prometheus) and samples were reduced with 4x NuPAGE LDS Sample Buffer (Life Technologies) containing 10%β-Mercaptoethanol. Samples were analyzed via SDS-PAGE and immunoblotting as stated above.

### Immunoprecipitation and mass spectrometry

Cells cultured in growth medium were rinsed with PBS++ and lysed with cold 25 mM Tris-HCl, pH 7.4, 150 mM NaCl, 1% Triton X-100, 5 mM MgCl_2_ and 2x protease and cOmplete Protease Inhibitor (ThermoFisher). Lysates were needle passed through a 21G syringe 10 times and incubated on ice for 20 minutes prior to being centrifuged at 4°C for 10 minutes at 13000x*g*. Lysate volume and protein content was equalized using a BCA protein assay kit (Prometheus). Lysates were incubated for 2 hours with 2 μg of anti-Notch1, 2 μg of anti-EGFR, or 2 μg of anti-E-Cadherin antibodies at 4 °C with rotation. Pierce Protein A/G beads (ThermoFisher) were equilibrated in cold lysis buffer prior to incubation with antibody complexes for 2 hours at 4 °C with rotation. Bead pellets were rinsed three times with cold lysis buffer and reduced with 1x NuPAGE LDS Sample Buffer (Life Technologies) containing 5% β-Mercaptoethanol. Samples were analyzed via SDS-PAGE and western blot.

Single, excised Coomassie-stained bands for protein identification were analyzed by MS Bioworks as follows. In-gel digestion was performed using a ProGest robot (DigiLab). Gel bands were washed with 25 mM ammonium bicarbonate followed by acetonitrile, reduced with 10 mM dithiothreitol at 60 °C followed by alkylation with 50 mM iodoacetamide at room temperature, digested with trypsin (Promega) at 37 °C f or 4 h, and quenched with formic acid and the supernatant was analyzed directly without further processing. Half of each digested sample was analyzed by nano LC-MS/MS with a Waters M-Class HPLC system interfaced to a ThermoFisher Fusion Lumos mass spectrometer. Peptides were loaded on a trapping column and eluted over a 75μm analytical column at 350nL/min; both columns were packed with Luna C18 resin (Phenomenex). The mass spectrometer was operated in data-dependent mode, with the Orbitrap operating at 60,000 FWHM and 15,000 FWHM for MS and MS/MS respectively.

### Statistical analysis

Sample sizes and P values are reported in each of the corresponding figure legends. Statistical analyses were performed in GraphPad Prism 8. Unless otherwise noted, graphs show mean ± standard error of the mean. When experiments involved only a single pair of conditions, statistical differences between the two sets of data were analyzed with a two-tailed, unpaired Student t-test assuming unequal variances. For data sets containing more than two samples, one-way ANOVA with a classical Bonferroni multiple-comparison post-test was used to determine adjusted P values. Images are representative of at least three independent experiments. Experiments were not randomized and the investigators were not blinded during data analysis. Source data for all graphs is provides in the Source Data Table.

### Materials and data availability

All unique reagents and data generated in this study are available from the lead contact without restriction.

## REFERENCES

Antfolk, D., Antila, C., Kemppainen, K., Landor, S.K., and Sahlgren, C. (2019). Decoding the PTM-switchboard of Notch. Biochim Biophys Acta Mol Cell Res 1866, 118507.

Belardi, B., Son, S., Felce, J.H., Dustin, M.L., and Fletcher, D.A. (2020). Cell-cell interfaces as specialized compartments directing cell function. Nat Rev Mol Cell Biol 21, 750–764.

Bentley, K., Franco, C.A., Philippides, A., Blanco, R., Dierkes, M., Gebala, V., Stanchi, F., Jones, M., Aspalter, I.M., Cagna, G., et al. (2014). The role of differential VE-cadherin dynamics in cell rearrangement during angiogenesis. Nat Cell Biol 16, 309–321.

Blanpain, C., Lowry, W.E., Pasolli, H.A., and Fuchs, E. (2006). Canonical notch signaling functions as a commitment switch in the epidermal lineage. Genes Dev 20, 3022–3035.

Borggrefe, T., and Oswald, F. (2009). The Notch signaling pathway: transcriptional regulation at Notch target genes. Cell Mol Life Sci 66, 1631–1646.

Borghi, N., and Nelson, W. (2009). Intercellular adhesion in morphogenesis: molecular and biophysical considerations. Curr Top Dev Biol 89, 1–32.

Bray, S.J. (2016). Notch signalling in context. Nat Rev Mol Cell Biol 17, 722–735.

Chiang, M.Y., Xu, M.L., Histen, G., Shestova, O., Roy, M., Nam, Y., Blacklow, S.C., Sacks, D.B., Pear, W.S., and Aster, J.C. (2006). Identification of a conserved negative regulatory sequence that influences the leukemogenic activity of NOTCH1. Mol Cell Biol 26, 6261–6271.

Chiasson-MacKenzie, C., Morris, Z.S., Baca, Q., Morris, B., Coker, J.K., Mirchev, R., Jensen, A.E., Carey, T., Stott, S.L., Golan, D.E., et al. (2015). NF2/Merlin mediates contact-dependent inhibition of EGFR mobility and internalization via cortical actomyosin. J Cell Biol 211, 391–405.

Chopra, A., Kutys, M.L., Zhang, K., Polacheck, W.J., Sheng, C.C., Luu, R.J., Eyckmans, J., Hinson, J.T., Seidman, J.G., Seidman, C.E., et al. (2018). Force Generation via beta-Cardiac Myosin, Titin, and alpha-Actinin Drives Cardiac Sarcomere Assembly from Cell-Matrix Adhesions. Dev Cell 44, 87–96 e85.

Collinet, C., and Lecuit, T. (2021). Programmed and self-organized flow of information during morphogenesis. Nat Rev Mol Cell Biol 22, 245–265.

Coon, B.G., Baeyens, N., Han, J., Budatha, M., Ross, T.D., Fang, J.S., Yun, S., Thomas, J.L., and Schwartz, M.A. (2015). Intramembrane binding of VE-cadherin to VEGFR2 and VEGFR3 assembles the endothelial mechanosensory complex. J Cell Biol 208, 975–986.

Crowner, D., Le Gall, M., Gates, M.A., and Giniger, E. (2003). Notch steers Drosophila ISNb motor axons by regulating the Abl signaling pathway. Curr Biol 13, 967–972.

Dahan, S., Rabinowitz, K.M., Martin, A.P., Berin, M.C., Unkeless, J.C., and Mayer, L. (2011). Notch-1 signaling regulates intestinal epithelial barrier function, through interaction with CD4+ T cells, in mice and humans. Gastroenterology 140, 550–559.

de Celis, J.F., Garcia-Bellido, A., and Bray, S.J. (1996). Activation and function of Notch at the dorsal-ventral boundary of the wing imaginal disc. Development 122, 359–369.

Demehri, S., Liu, Z., Lee, J., Lin, M.H., Crosby, S.D., Roberts, C.J., Grigsby, P.W., Miner, J.H., Farr, A.G., and Kopan, R. (2008). Notch-deficient skin induces a lethal systemic B-lymphoproliferative disorder by secreting TSLP, a sentinel for epidermal integrity. PLoS Biol 6, e123.

Demehri, S., Turkoz, A., and Kopan, R. (2009). Epidermal Notch1 loss promotes skin tumorigenesis by impacting the stromal microenvironment. Cancer Cell 16, 55–66.

Diaz-Rohrer, B.B., Levental, K.R., Simons, K., and Levental, I. (2014). Membrane raft association is a determinant of plasma membrane localization. Proc Natl Acad Sci U S A 111, 8500–8505.

Ding, Y., Estrella, M.R., Hu, Y.Y., Chan, H.L., Zhang, H.D., Kim, J.W., Simmer, J.P., and Hu, J.C. (2009). Fam83h is associated with intracellular vesicles and ADHCAI. J Dent Res 88, 991–996.

Falo-Sanjuan, J., and Bray, S. (2022). Notch-dependent and -independent transcription are modulated by tissue movements at gastrulation. Elife 11.

Falo-Sanjuan, J., and Bray, S.J. (2021). Membrane architecture and adherens junctions contribute to strong Notch pathway activation. Development 148.

Gordon, W.R., Arnett, K.L., and Blacklow, S.C. (2008). The molecular logic of Notch signaling--a structural and biochemical perspective. J Cell Sci 121, 3109–3119.

Gordon, W.R., Zimmerman, B., He, L., Miles, L.J., Huang, J., Tiyanont, K., McArthur, D.G., Aster, J.C., Perrimon, N., Loparo, J.J., et al. (2015). Mechanical Allostery: Evidence for a Force Requirement in the Proteolytic Activation of Notch. Dev Cell 33, 729–736.

Grammont, M. (2007). Adherens junction remodeling by the Notch pathway in Drosophila melanogaster oogenesis. J Cell Biol 177, 139–150.

Gumbiner, B.M. (1996). Cell adhesion: the molecular basis of tissue architecture and morphogenesis. Cell 84, 345–357.

Guo, Z., Neilson, L.J., Zhong, H., Murray, P.S., Zanivan, S., and Zaidel-Bar, R. (2014). E-cadherin interactome complexity and robustness resolved by quantitative proteomics. Sci Signal 7, rs7.

Hansson, E.M., Teixeira, A.I., Gustafsson, M.V., Dohda, T., Chapman, G., Meletis, K., Muhr, J., and Lendahl, U. (2006). Recording Notch signaling in real time. Dev Neurosci 28, 118–127.

Hellstrom, M., Phng, L.K., Hofmann, J.J., Wallgard, E., Coultas, L., Lindblom, P., Alva, J., Nilsson, A.K., Karlsson, L., Gaiano, N., et al. (2007). Dll4 signalling through Notch1 regulates formation of tip cells during angiogenesis. Nature 445, 776–780.

Hunter, G.L., He, L., Perrimon, N., Charras, G., Giniger, E., and Baum, B. (2019). A role for actomyosin contractility in Notch signaling. BMC Biol 17, 12.

Jaffe, A.B., Kaji, N., Durgan, J., and Hall, A. (2008). Cdc42 controls spindle orientation to position the apical surface during epithelial morphogenesis. J Cell Biol 183, 625–633.

Jheon, A.H., Prochazkova, M., Meng, B., Wen, T., Lim, Y.J., Naveau, A., Espinoza, R., Cox, T.C., Sone, E.D., Ganss, B., et al. (2016). Inhibition of Notch Signaling During Mouse Incisor Renewal Leads to Enamel Defects. J Bone Miner Res 31, 152–162.

Johnson, J.L., Najor, N.A., and Green, K.J. (2014). Desmosomes: regulators of cellular signaling and adhesion in epidermal health and disease. Cold Spring Harb Perspect Med 4, a015297.

Khait, I., Orsher, Y., Golan, O., Binshtok, U., Gordon-Bar, N., Amir-Zilberstein, L., and Sprinzak, D. (2016). Quantitative Analysis of Delta-like 1 Membrane Dynamics Elucidates the Role of Contact Geometry on Notch Signaling. Cell Rep 14, 225–233.

Kim, J.W., Lee, S.K., Lee, Z.H., Park, J.C., Lee, K.E., Lee, M.H., Park, J.T., Seo, B.M., Hu, J.C., and Simmer, J.P. (2008). FAM83H mutations in families with autosomal-dominant hypocalcified amelogenesis imperfecta. Am J Hum Genet 82, 489–494.

Kim, K.M., Hussein, U.K., Park, S.H., Kang, M.A., Moon, Y.J., Zhang, Z., Song, Y., Park, H.S., Bae, J.S., Park, B.H., et al. (2019). FAM83H is involved in stabilization of beta-catenin and progression of osteosarcomas. J Exp Clin Cancer Res 38, 267.

Kopan, R., and Ilagan, M.X. (2009). The canonical Notch signaling pathway: unfolding the activation mechanism. Cell 137, 216–233.

Kovall, R.A., Gebelein, B., Sprinzak, D., and Kopan, R. (2017). The Canonical Notch Signaling Pathway: Structural and Biochemical Insights into Shape, Sugar, and Force. Dev Cell 41, 228–241.

Kuga, T., Sasaki, M., Mikami, T., Miake, Y., Adachi, J., Shimizu, M., Saito, Y., Koura, M., Takeda, Y., Matsuda, J., et al. (2016). FAM83H and casein kinase I regulate the organization of the keratin cytoskeleton and formation of desmosomes. Sci Rep 6, 26557.

Kutys, M.L., Polacheck, W.J., Welch, M.K., Gagnon, K.A., Koorman, T., Kim, S., Li, L., McClatchey, A.I., and Chen, C.S. (2020). Uncovering mutation-specific morphogenic phenotypes and paracrine-mediated vessel dysfunction in a biomimetic vascularized mammary duct platform. Nat Commun 11, 3377.

Kwak, M., Southard, K.M., Kim, W.R., Lin, A., Kim, N.H., Gopalappa, R., Lee, H.J., An, M., Choi, S.H., Jung, Y., et al. (2022). Adherens junctions organize size-selective proteolytic hotspots critical for Notch signalling. Nat Cell Biol 24, 1739–1753.

Lahdeniemi, I.A.K., Misiorek, J.O., Antila, C.J.M., Landor, S.K., Stenvall, C.A., Fortelius, L.E., Bergstrom, L.K., Sahlgren, C., and Toivola, D.M. (2017). Keratins regulate colonic epithelial cell differentiation through the Notch1 signalling pathway. Cell Death Differ 24, 984–996.

Lemmon, M.A., and Schlessinger, J. (2010). Cell signaling by receptor tyrosine kinases. Cell 141, 1117–1134.

Lewis, J.D., Caldara, A.L., Zimmer, S.E., Stahley, S.N., Seybold, A., Strong, N.L., Frangakis, A.S., Levental, I., Wahl, J.K., 3rd, Mattheyses, A.L., et al. (2019). The desmosome is a mesoscale lipid raft-like membrane domain. Mol Biol Cell 30, 1390–1405.

Lisica, A., Fouchard, J., Kelkar, M., Wyatt, T.P.J., Duque, J., Ndiaye, A.B., Bonfanti, A., Baum, B., Kabla, A.J., and Charras, G.T. (2022). Tension at intercellular junctions is necessary for accurate orientation of cell division in the epithelium plane. Proc Natl Acad Sci U S A 119, e2201600119.

Lloyd-Lewis, B., Mourikis, P., and Fre, S. (2019). Notch signalling: sensor and instructor of the microenvironment to coordinate cell fate and organ morphogenesis. Curr Opin Cell Biol 61, 16–23.

Lowell, S., and Watt, F.M. (2001). Delta regulates keratinocyte spreading and motility independently of differentiation. Mech Dev 107, 133–140.

Major, R.J., and Irvine, K.D. (2005). Influence of Notch on dorsoventral compartmentalization and actin organization in the Drosophila wing. Development 132, 3823–3833.

Major, R.J., and Irvine, K.D. (2006). Localization and requirement for Myosin II at the dorsal-ventral compartment boundary of the Drosophila wing. Dev Dyn 235, 3051–3058.

Mertz, A.F., Che, Y., Banerjee, S., Goldstein, J.M., Rosowski, K.A., Revilla, S.F., Niessen, C.M., Marchetti, M.C., Dufresne, E.R., and Horsley, V. (2013). Cadherin-based intercellular adhesions organize epithelial cell-matrix traction forces. Proc Natl Acad Sci U S A 110, 842–847.

Movahedan, A., Afsharkhamseh, N., Sagha, H.M., Shah, J.R., Milani, B.Y., Milani, F.Y., Logothetis, H.D., Chan, C.C., and Djalilian, A.R. (2013). Loss of Notch1 disrupts the barrier repair in the corneal epithelium. PLoS One 8, e69113.

Nicolas, M., Wolfer, A., Raj, K., Kummer, J.A., Mill, P., van Noort, M., Hui, C.C., Clevers, H., Dotto, G. P., and Radtke, F. (2003). Notch1 functions as a tumor suppressor in mouse skin. Nat Genet 33, 416–421.

Nyga, A., Munoz, J.J., Dercksen, S., Fornabaio, G., Uroz, M., Trepat, X., Baum, B., Matthews, H. K., and Conte, V. (2021). Oncogenic RAS instructs morphological transformation of human epithelia via differential tissue mechanics. Sci Adv 7, eabg6467.

Oldenburg, J., van der Krogt, G., Twiss, F., Bongaarts, A., Habani, Y., Slotman, J.A., Houtsmuller, A., Huveneers, S., and de Rooij, J. (2015). VASP, zyxin and TES are tension-dependent members of Focal Adherens Junctions independent of the alpha-catenin-vinculin module. Sci Rep 5, 17225.

Polacheck, W.J., Kutys, M.L., Tefft, J.B., and Chen, C.S. (2019). Microfabricated blood vessels for modeling the vascular transport barrier. Nat Protoc 14, 1425–1454.

Polacheck, W.J., Kutys, M.L., Yang, J., Eyckmans, J., Wu, Y., Vasavada, H., Hirschi, K.K., and Chen, C.S. (2017). A non-canonical Notch complex regulates adherens junctions and vascular barrier function. Nature 552, 258–262.

Prechova, M., Adamova, Z., Schweizer, A.L., Maninova, M., Bauer, A., Kah, D., Meier-Menches, S.M., Wiche, G., Fabry, B., and Gregor, M. (2022). Plectin-mediated cytoskeletal crosstalk controls cell tension and cohesion in epithelial sheets. J Cell Biol 221.

Priya, R., Allanki, S., Gentile, A., Mansingh, S., Uribe, V., Maischein, H.M., and Stainier, D.Y.R. (2020). Tension heterogeneity directs form and fate to pattern the myocardial wall. Nature 588, 130–134.

Qian, X., Karpova, T., Sheppard, A.M., McNally, J., and Lowy, D.R. (2004). E-cadherin-mediated adhesion inhibits ligand-dependent activation of diverse receptor tyrosine kinases. EMBO J 23, 1739–1748.

Scarpa, E., Szabo, A., Bibonne, A., Theveneau, E., Parsons, M., and Mayor, R. (2015). Cadherin Switch during EMT in Neural Crest Cells Leads to Contact Inhibition of Locomotion via Repolarization of Forces. Dev Cell 34, 421–434.

Shaya, O., Binshtok, U., Hersch, M., Rivkin, D., Weinreb, S., Amir-Zilberstein, L., Khamaisi, B., Oppenheim, O., Desai, R.A., Goodyear, R.J., et al. (2017). Cell-Cell Contact Area Affects Notch Signaling and Notch-Dependent Patterning. Dev Cell 40, 505–511 e506.

Siebel, C., and Lendahl, U. (2017). Notch Signaling in Development, Tissue Homeostasis, and Disease. Physiol Rev 97, 1235–1294.

Snijders, A.M., Lee, S.Y., Hang, B., Hao, W., Bissell, M.J., and Mao, J.H. (2017). FAM83 family oncogenes are broadly involved in human cancers: an integrative multi-omics approach. Mol Oncol 11, 167–179.

Sullivan, B., Light, T., Vu, V., Kapustka, A., Hristova, K., and Leckband, D. (2022). Mechanical disruption of E-cadherin complexes with epidermal growth factor receptor actuates growth factor-dependent signaling. Proc Natl Acad Sci U S A 119.

Tokuchi, K., Kitamura, S., Maeda, T., Watanabe, M., Hatakeyama, S., Kano, S., Tanaka, S., Ujiie, H., and Yanagi, T. (2021). Loss of FAM83H promotes cell migration and invasion in cutaneous squamous cell carcinoma via impaired keratin distribution. J Dermatol Sci 104, 112–121.

Totaro, A., Castellan, M., Battilana, G., Zanconato, F., Azzolin, L., Giulitti, S., Cordenonsi, M., and Piccolo, S. (2017). YAP/TAZ link cell mechanics to Notch signalling to control epidermal stem cell fate. Nat Commun 8, 15206.

Tseng, Q., Duchemin-Pelletier, E., Deshiere, A., Balland, M., Guillou, H., Filhol, O., and Thery, M. (2012). Spatial organization of the extracellular matrix regulates cell-cell junction positioning. Proc Natl Acad Sci U S A 109, 1506–1511.

Wang, H., Zang, C., Taing, L., Arnett, K.L., Wong, Y.J., Pear, W.S., Blacklow, S.C., Liu, X.S., and Aster, J.C. (2014). NOTCH1-RBPJ complexes drive target gene expression through dynamic interactions with superenhancers. Proc Natl Acad Sci U S A 111, 705–710.

Wang, S.K., Hu, Y., Yang, J., Smith, C.E., Richardson, A.S., Yamakoshi, Y., Lee, Y.L., Seymen, F., Koruyucu, M., Gencay, K., et al. (2016). Fam83h null mice support a neomorphic mechanism for human ADHCAI. Mol Genet Genomic Med 4, 46–67.

Weng, A.P., Nam, Y., Wolfe, M.S., Pear, W.S., Griffin, J.D., Blacklow, S.C., and Aster, J.C. (2003). Growth suppression of pre-T acute lymphoblastic leukemia cells by inhibition of notch signaling. Mol Cell Biol 23, 655–664.

Wu, J., Tannan, N.B., Vuong, L.T., Koca, Y., Collu, G.M., and Mlodzik, M. (2022). Par3/Bazooka promotes Notch pathway target gene activation. 2022.2005.2024.493322.

Zakirov, B., Charalambous, G., Thuret, R., Aspalter, I.M., Van-Vuuren, K., Mead, T., Harrington, K., Regan, E.R., Herbert, S.P., and Bentley, K. (2021). Active perception during angiogenesis: filopodia speed up Notch selection of tip cells in silico and in vivo. Philos Trans R Soc Lond B Biol Sci 376, 20190753.

