## Supplementary Information for "Notch1 cortical signaling regulates epithelial architecture and cell-cell adhesion"

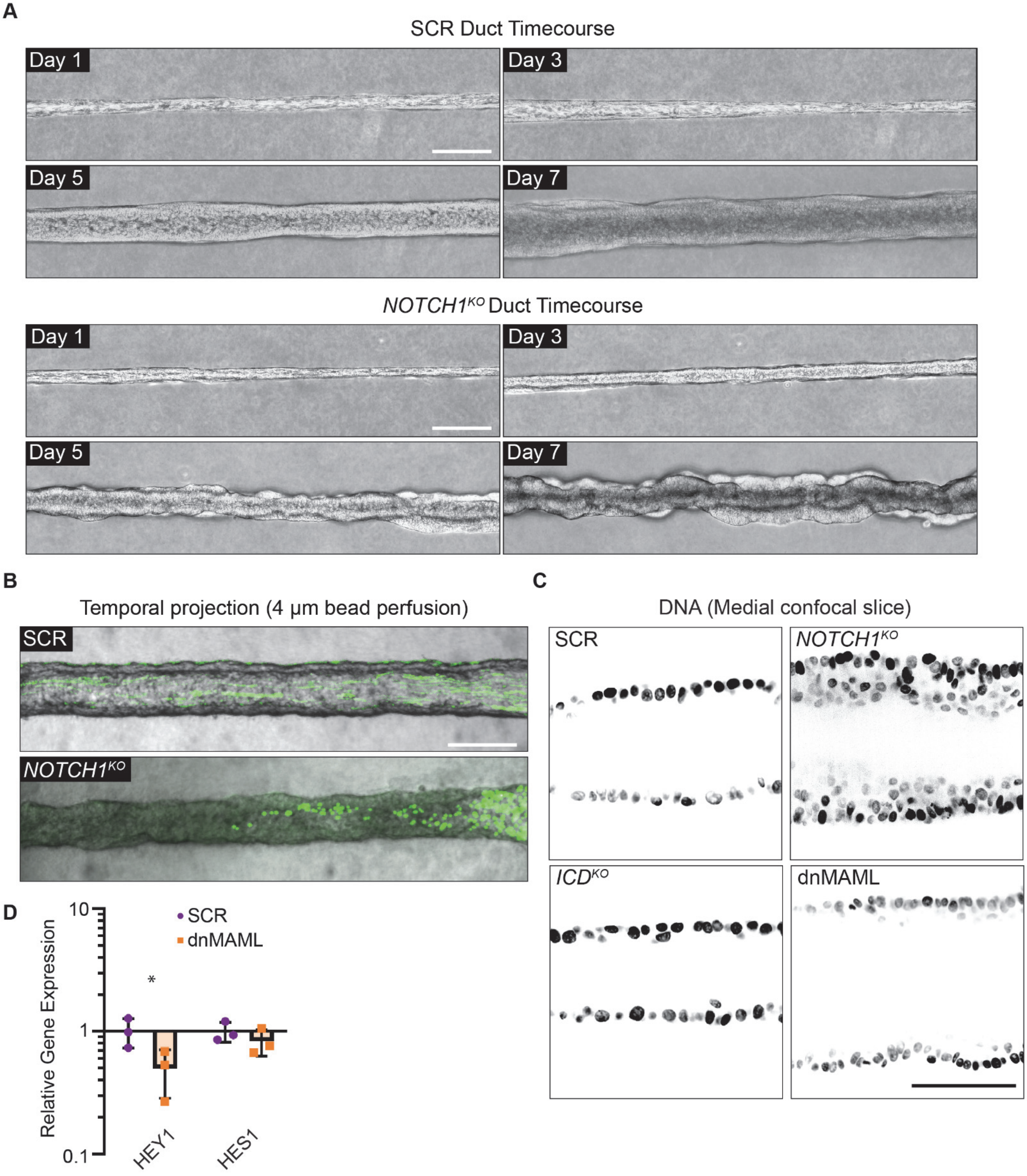


**Supplemental Figure 1. Notch1 cortical signaling influences mammary duct morphogenesis.** (A) Top: Phase contrast micrographs of scramble control (SCR) mammary ducts over a 7-day timecourse shown at days 1, 3, 5, and 7 post-seeding. Bottom: Phase-contrast images of *NOTCH1$^{KO}$* mammary ducts over a 7-day timecourse shown at days 1, 3, 5, and 7 post-seeding. Scale bars, 150 µm. (B) Temporal projection micrographs of a timelapse of SCR and *NOTCH1$^{KO}$* mammary epithelial ducts perfused with 4 µm polystyrene beads (green). Scale bar, 200 µm. (C) Medial confocal slice micrographs from SCR, *NOTCH1$^{KO}$*, *ICD$^{KO}$*, and dnMAML labeled with Hoechst (black). Scale bar, 50 µm. (D) Expression of Notch1-target genes HES1 and HEY1 measured by qPCR in SCR or dnMAML-expressing monolayers. n = 3 independent experiments. For plot D, mean ± SEM; two-tailed unpaired t test, *$p<0.05$.

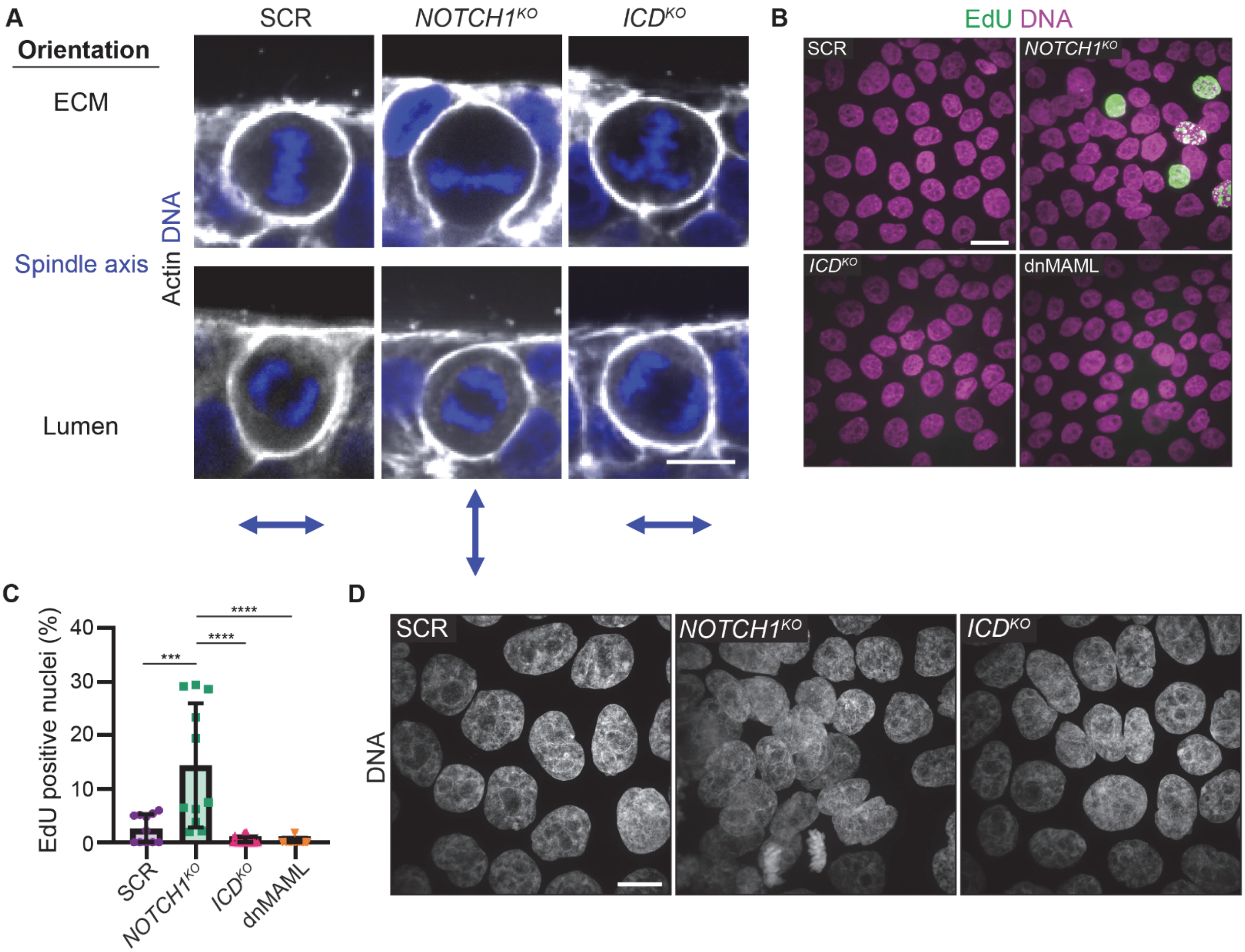


**Supplemental Figure 2. Loss of Notch1 cortical signaling results in aberrant epithelial architecture and proliferation.** (A) Fluorescence micrographs of two sets of dividing scramble control (SCR), $NOTCH1^{KO}$, and $ICD^{KO}$ cells in ducts labeled with Hoechst (blue) and phalloidin (grey). Scale bar, 10 µm. Top and bottom rows are independent cells, during metaphase or anaphase respectively. (B) Fluorescence micrographs of SCR, $NOTCH1^{KO}$, $ICD^{KO}$, and dnMAML monolayers labeled with EdU (green) and Hoechst (magenta). Scale bar, 20 µm. (C) Quantification of the percentage of EdU positive nuclei in SCR, $NOTCH1^{KO}$, $ICD^{KO}$, and dnMAML monolayers. n ≥ 12 fields of view, from three independent experiments. (D) Maximum intensity projection fluorescence micrographs of SCR, $NOTCH1^{KO}$, and $ICD^{KO}$ monolayers labeled with Hoechst (grey). Scale bar, 10 µm. For plot C, mean ± SEM; one-way ANOVA with Tukey's post-hoc test, ***$p<0.001$, ****$p<0.0001$.

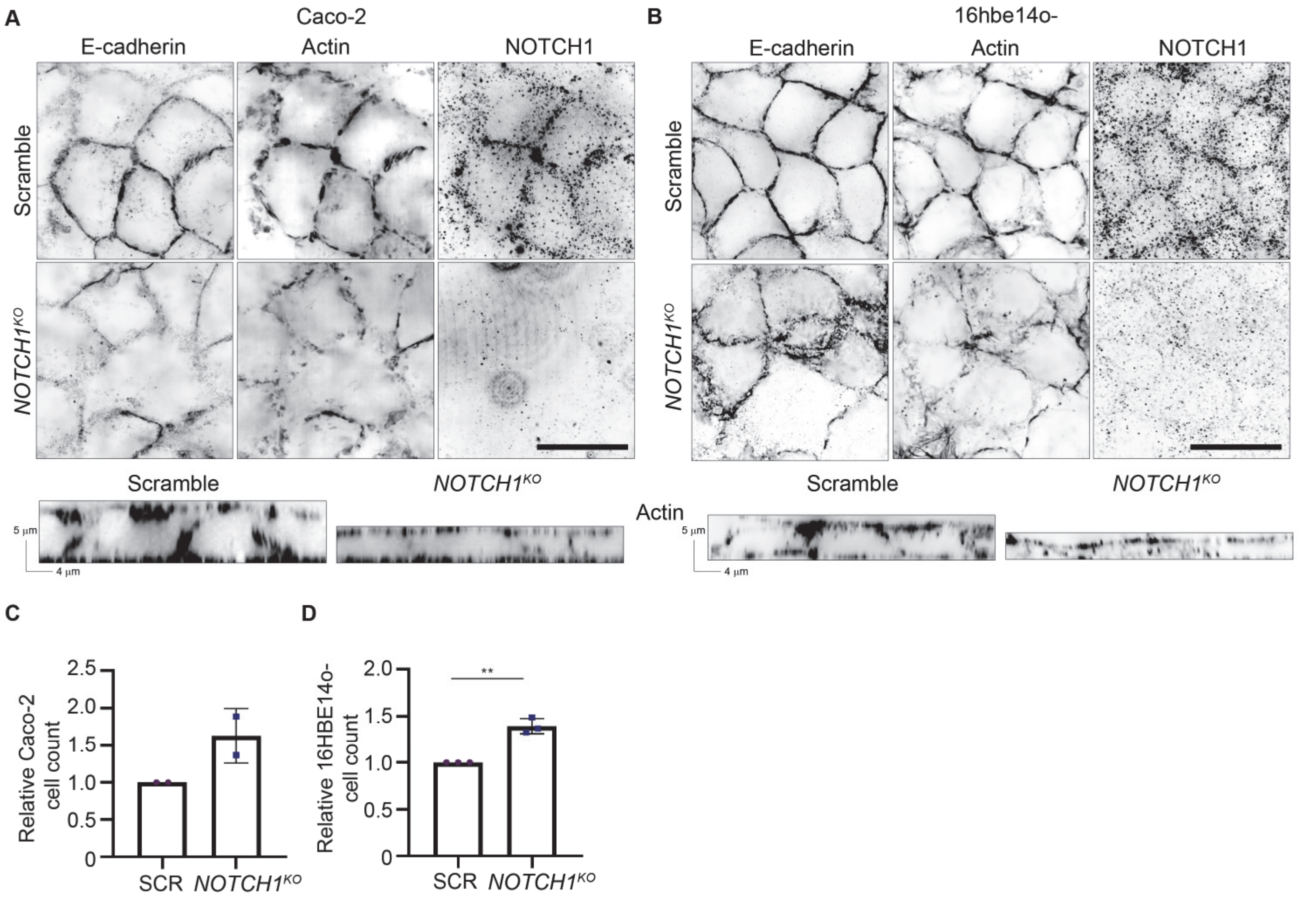


**Supplemental Figure 3. Notch1 regulates epithelial architecture, adherens junctions, cortical actin organization, and proliferation in human intestinal and bronchial epithelia.** (A) Top: Immunofluorescence micrographs of scramble control (SCR, top row) and *NOTCH1*$^{KO}$ (bottom row) Caco-2 monolayers immunostained for E-Cadherin (black) and Notch1 (black) and labeled with phalloidin (black). Bottom: YZ orthogonal projections from micrographs of SCR and *NOTCH1*$^{KO}$ Caco-2 monolayers labeled with phalloidin (black). Scale bar, 20 µm. (B) Top: Fluorescence micrographs of SCR (top row) and *NOTCH1*$^{KO}$ (bottom row) 16hbe14o- monolayers immunostained for E-Cadherin (black) and Notch1 (black) and labeled with phalloidin (black). Bottom: YZ orthogonal projections from micrographs of SCR and *NOTCH1*$^{KO}$ 16hbe14o- monolayers labeled with phalloidin (black). Scale bar, 20 µm. (C) Relative number of SCR and *NOTCH1*$^{KO}$ Caco-2 cells measured at passage. n = 2 plates from two independent experiments. (D) Relative number of SCR and *NOTCH1*$^{KO}$ 16hbe14o- cells measured at passage. *n* = 3 plates from three independent experiments. For plot D, mean ± SEM; two-tailed unpaired t test, **p<0.01.

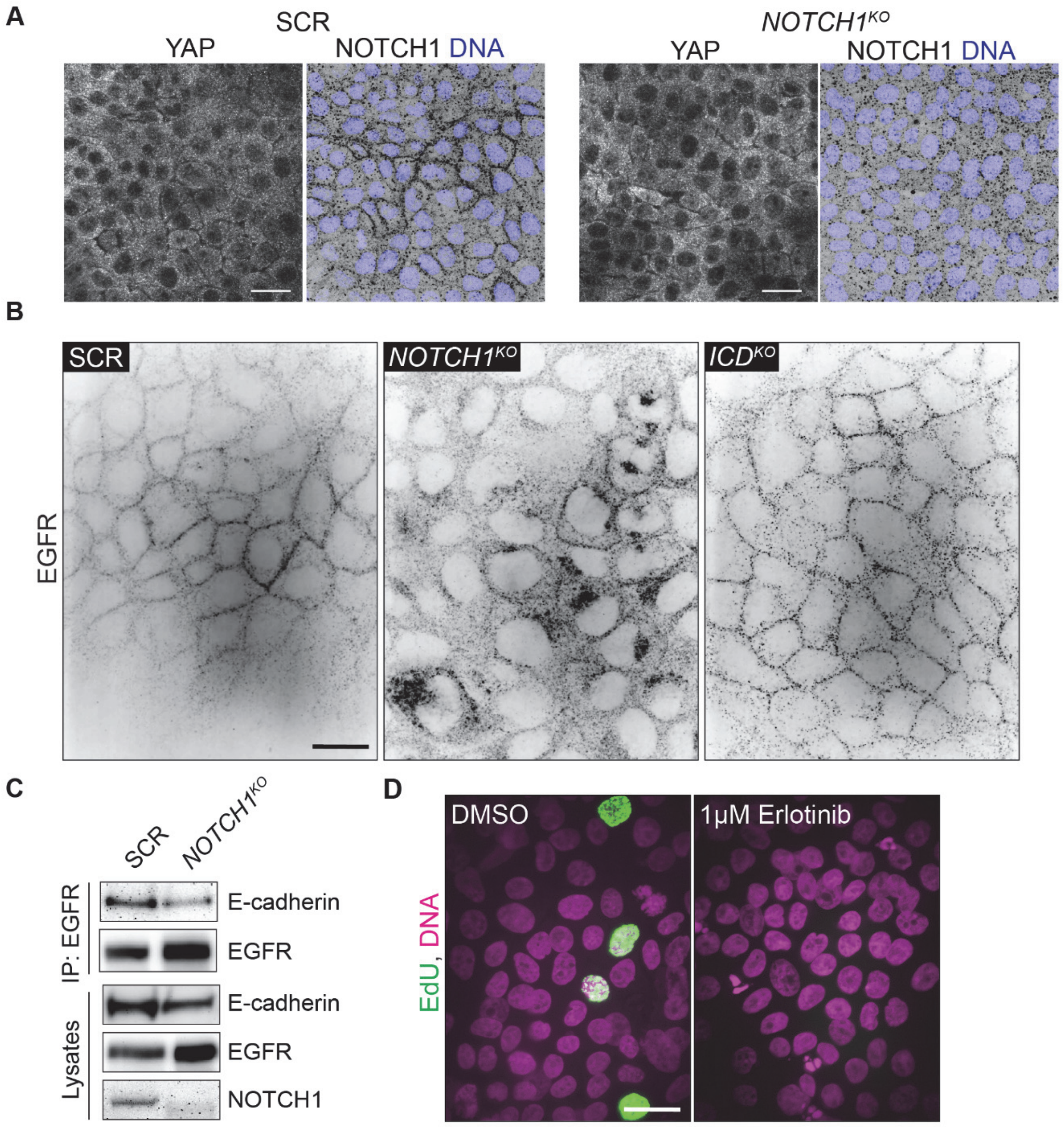


**Supplemental Figure 4. Mitogenic signaling in *NOTCH1*$^{KO}$.** (A) Immunofluorescence micrographs of scramble control (SCR) and *NOTCH1*$^{KO}$ monolayers immunostained for YAP (white, left) and Notch1 (black) and labeled with DAPI (blue) (right). Scale bar, 20 μm. (B) Full field of view source fluorescence micrographs of SCR, *NOTCH1*$^{KO}$, and *ICD*$^{KO}$ monolayers immunostained for EGFR (black) in Figure 2. Scale bar, 20 μm. (C) Western blot of immunoprecipitation of EGFR from SCR and *NOTCH1*$^{KO}$ monolayer lysates immunoblotted for E-Cadherin and EGFR. Representative of three independent experiments. (D) Fluorescence micrographs of *NOTCH1*$^{KO}$ monolayers treated with DMSO or 1 μM Erlotinib labeled with EdU (green) and Hoechst (pink). Scale bar, 20 μm.

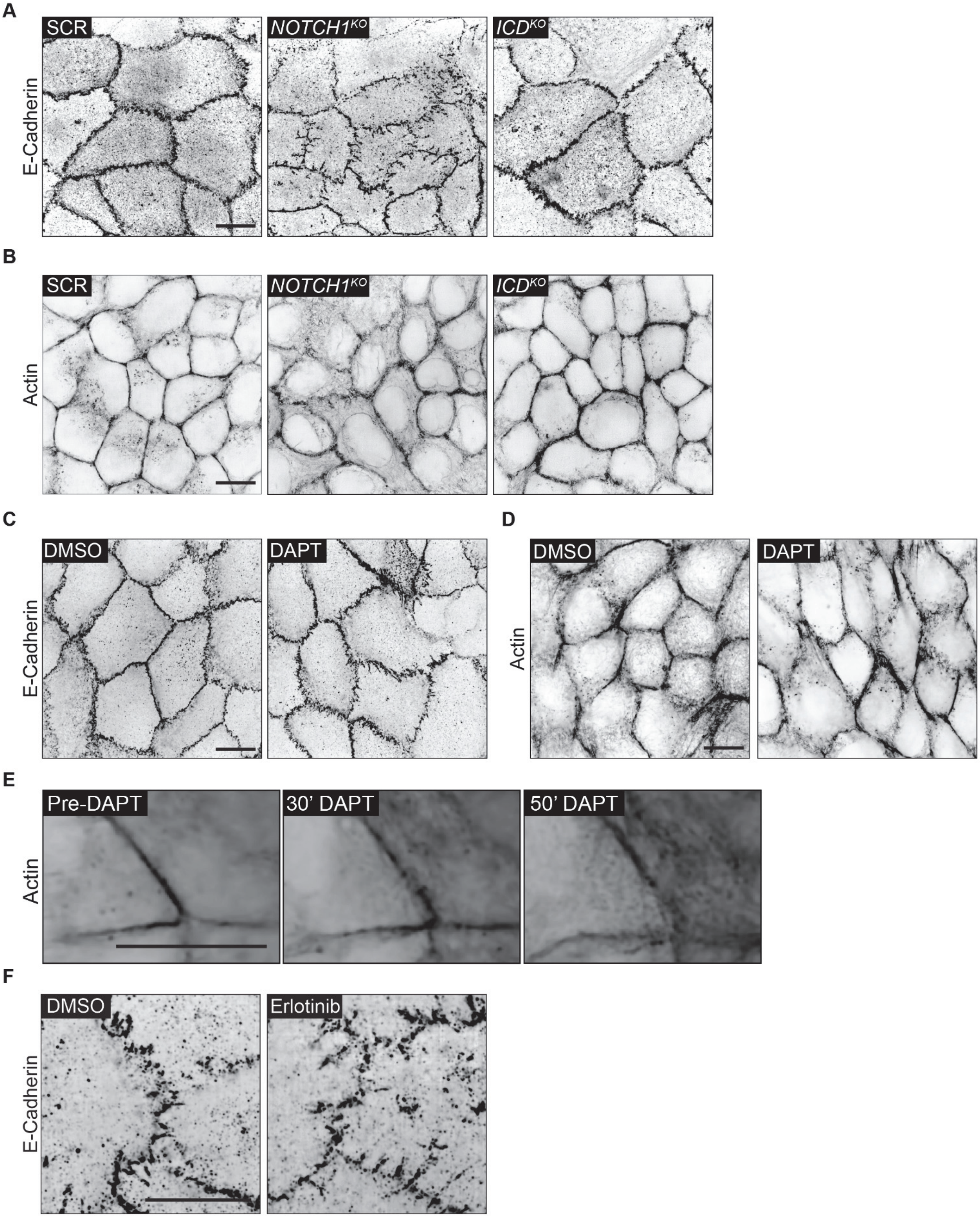


**Supplemental Figure 5. Loss of Notch1 cortical signaling disrupts epithelial adherens junctions and cortical actin organization.** (A) Source full field of view fluorescence micrographs of SCR, $NOTCH1^{KO}$, and $ICD^{KO}$ monolayers immunostained for E-Cadherin (black) for representative images in Figure 3. (B) Source full field of view fluorescence micrographs of SCR, $NOTCH1^{KO}$, and $ICD^{KO}$ monolayers labeled with phalloidin (black) for representative images in Figure 3. (C) Source full field of view fluorescence micrographs of wild type cells treated with DMSO or 10 μM DAPT for two hours immunostained for E-Cadherin (black) and (D) actin (black) for representative images in Figure 3. (E) Fluorescence micrographs of single frames from a timelapse movie of wild-type cells labeled with SPY650-FastAct (black) and treated with 10 μM DAPT. (F) Immunofluorescence micrographs of $NOTCH1^{KO}$ monolayers treated with DMSO or 1 μM Erlotinib immunostained for E-Cadherin (black). All scale bars, 10 μm.

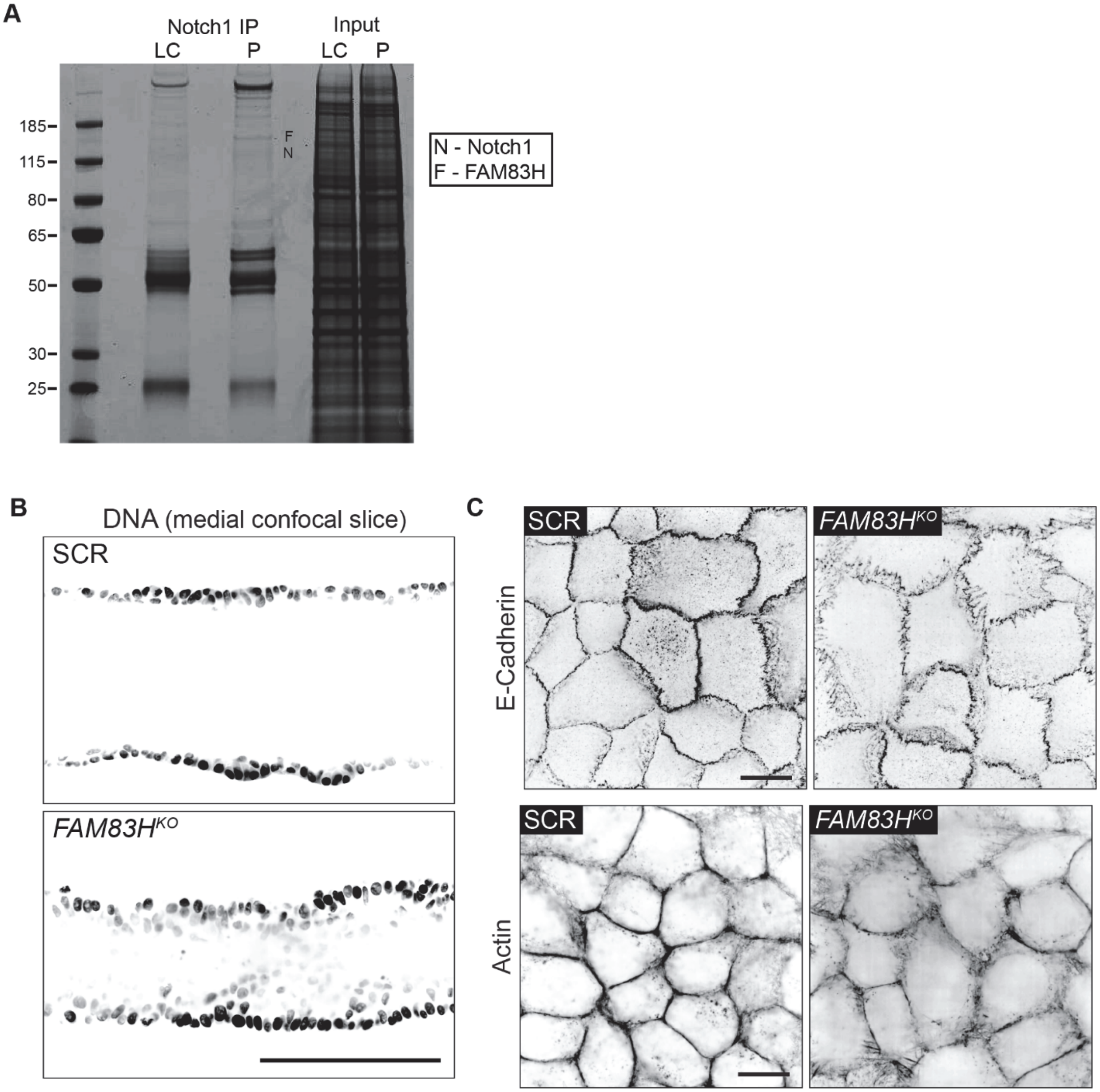


**Supplemental Figure 6. Loss of FAM83H leads to focal adherens junctions, depleted cortical actin, and alterations in ductal architecture.** (A) Representative Coomassie stained SDS-PAGE gel of Notch1 immunoprecipitation from LC and P monolayers. F – denotes the band identified as FAM83H by mass spectrometry. (B) Medial confocal slices micrographs of scramble control (SCR) and *FAM83H*$^{KO}$ labeled with Hoechst (black). Scale bar, 100 µm. (C) Immunofluorescence micrographs of SCR and *FAM83H*$^{KO}$ monolayers stained with E-Cadherin and labeled with phalloidin. Scale bars, 10 µm.

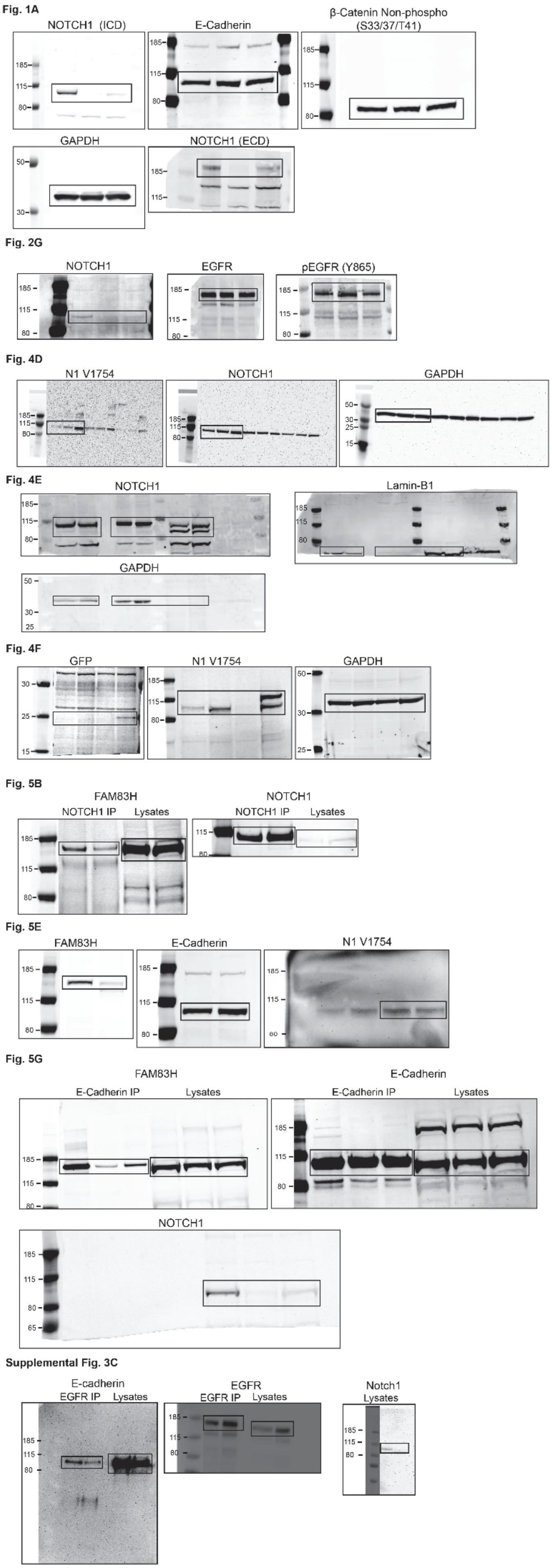


**Supplemental Figure 7.** Uncropped western blot images.

**Supplementary Video 1.** Live cell imaging of the medial confocal plane of a SPY650-FastAct labeled *NOTCH1*$^{KO}$ duct during assembly. Asterisks denote representative cell divisions. Scale bar, 100 μm. Time scale, hour:minute, displayed at 6000x speed.
